# Significant abundance of *cis* configurations of mutations in diploid human genomes

**DOI:** 10.1101/221085

**Authors:** Margret R. Hoehe, Ralf Herwig, Qing Mao, Brock A. Peters, Radoje Drmanac, George M. Church, Thomas Huebsch

## Abstract

To fully understand human genetic variation, one must assess the specific distribution of variants between the two chromosomal homologues of genes, and any functional units of interest, as the phase of variants can significantly impact gene function and phenotype. To this end, we have systematically analyzed 18,121 autosomal protein-coding genes in 1,092 statistically phased genomes from the 1000 Genomes Project, and an unprecedented number of 184 experimentally phased genomes from the Personal Genome Project. Here we show that mutations predicted to functionally alter the protein, and coding variants as a whole, are not randomly distributed between the two homologues of a gene, but do occur significantly more frequently in *cis*-than *trans*-configurations, with *cis/trans* ratios of ∼60:40. Significant *cis*-abundance was observed in virtually all individual genomes in all populations. Nearly all variable genes exhibited either *cis*, or *trans* configurations of protein-altering mutations in significant excess, allowing distinction of *cis*- and *trans*-abundant genes. These common patterns of phase were largely constituted by a shared, global set of phase-sensitive genes. We show significant enrichment of this global set with gene sets indicating its involvement in adaptation and evolution. Moreover, *cis*- and *trans*-abundant genes were found functionally distinguishable, and exhibited strikingly different distributional patterns of protein-altering mutations. This work establishes common patterns of phase as key characteristics of diploid human exomes and provides evidence for their potential functional significance. Thus, it highlights the importance of phase for the interpretation of protein-coding genetic variation, challenging the current conceptual and functional interpretation of autosomal genes.

## Introduction

The analysis of human genetic variation has focused so far on the identification, cataloguing and annotation of variants, particularly in protein-coding genes (Abecasis et al. 2012; Lek et al. 2016). One key aspect of genetic variation has, however, not yet been fully addressed: the distribution of variants between the two homologous chromosomes in a diploid genome. Whether variants reside on the same homologue, in *cis*, or on both homologues, in *trans*, is “key to understanding their impact on gene function and phenotype” (Suk et al. 2011; Tewhey et al. 2011). For example, while two null mutations in *cis* leave the second form of the gene intact, no functional form of the gene is present in a *trans* configuration, changing phenotype entirely (Benzer 1957). Thus, to fully understand the biology of genes and genomes, the phase of mutations must be known. In a first population level analysis of haplotype-resolved genomes (Hoehe et al. 2014), we have shown non-random distribution of mutations between homologous gene forms, as indicated by significant abundance of *cis* configurations with an average *cis/trans* ratio of approximately 60:40. This raises the question: Could *cis*- abundance of mutations, the preferential location of mutations on the same chromosomal homologue of an autosomal gene, represent a key characteristic of diploid human genomes? And if so, which types of genes constitute such phase imbalance? Thus, we aimed to (i) assess the distribution of mutations between protein-coding homologous gene forms in substantially larger numbers of genomes and different population samples, (ii) determine, significant abundance of *cis* configurations given, the types of genes underlying this phenomenon and (iii) uncover potential biological implications of phase imbalance.

We assessed the *cis/trans* ratios of mutations in 1,092 statistically haplotype-resolved genomes from the 1000 Genomes database (Abecasis et al. 2012) and each of the different populations contained therein separately. Because the analysis of molecularly and statistically haplotype-resolved genomes had yielded nearly identical results earlier (Hoehe et al. 2014), the use of this large resource appeared suitable. The key results obtained from this resource were then cross-validated in an independent sample of 184 experimentally phased genomes from the Personal Genome Project (PGP) (Mao et al. 2016; Ball et al. 2012), a huge advance over the recent population-level analysis of 14 molecularly haplotype-resolved genomes (Hoehe et al. 2014). *Cis/trans* ratios were assessed for population samples as a whole as well as for each individual genome separately as described in more detail in Box 1, which also provides further information on the concepts underlying our approach. The ratios were determined for mutations predicted to alter protein function (the term “protein-altering” is used synonymously in text), the entirety of amino acid (AA) exchanges and synonymous SNPs (sSNPs), and the combinations of these as they co-occur within the genes. Significant abundance of *cis* configurations evident, the *cis/trans* ratios were dissected further per number of variants in a configuration, and the observed ratios compared to the theoretically expected ratios to estimate *cis* excess. To explain this phenomenon, we examined evidence for ancestral admixture as a potential underlying mechanism. To address the functional implications of this phenomenon, we focused on the genes that had ≥ 2 mutations predicted to functionally alter the protein in either *cis*, or *trans* configurations. The fractions of these genes being highly similar between the genomes, we demonstrated existence of a shared, global set of phase-sensitive genes. Testing the genes in this set subsequently for significant excess of either *cis* or *trans* configurations unveiled a larger group of *cis*-and smaller group of *trans*-abundant genes, resulting in observed *cis/trans* ratio of about 60:40 as the net effect. These two categories of genes were further functionally differentiated by gene ontology (GO) and pathway analysis, and characterized by different mutational architectures of their phase configurations with potential implications for protein structure. Thus, the distinction of *cis* and *trans* configurations ultimately leads to the classification of variable autosomal genes into two major, potentially functionally divergent, categories. This work provides a first phase-informed global view of the diploid human exome, highlighting the importance of phase for the interpretation of protein-coding genetic variation, with implications for the conceptual and functional interpretation of autosomal genes.

#### BOX 1 *Cis* and *trans* configurations of mutations in protein-coding genes

##### Scheme 1 *Cis* and *trans* configuration of mutations

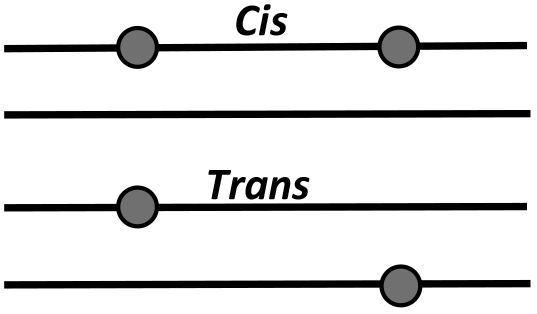

Where diploid autosomal genes have ≥ 2 variants, the distribution of these variants between the two chromosomal homologues, their phase, needs to be determined. In principle, the variants can be located on the same homologue, in a *cis* configuration, or on both homologues, in a *trans* configuration (Scheme 1). The genes with ≥ 2 variants, which could exist in either configuration, are also defined as ‘phase-sensitive’. Phase is determined (by convention) for nucleotides different from the reference sequence, usually the minor allele. The distinction of different phase configurations is biologically meaningful, where the variants are potentially functionally significant, such as mutations in the protein-coding sequence.

Mutation annotation is achieved by algorithmic prediction (see Methods). For the terms such as ‘potentially damaging’ or ‘damaging’ according to different programs, the term ‘protein-altering’ is used synonymously in text. The phase configurations of the totals of amino acid exchanges and synonymous SNPs, respectively, are examined in addition to serve as a control and allow differentiation of potentially underlying mechanisms of *cis*-abundance, i.e. selection of protein-altering mutations versus a common mechanism underlying all types of coding variants. For control, also the phase configurations including the entirety of these types of coding variants, as they co-occur within the genes, are assessed. The distinction of a *cis* or *trans* configuration requires haplotype information, which we obtained from the 1000G database (Abecasis et al. 2012) or the experimentally phased PGP genomes (Mao et al. 2016). Importantly, in the latter source, only those variants were included in analysis, which were contained within the same contig. Notably, in this work, we refer to the pair of haplotypes per gene per individual as the unit of analysis, which needs to be distinguished from the population-based (common) haplotypes inferred in the HapMap and 1000G. While our approach to the analysis of haplotypes concerns the potential biological implications of phase, the prevailing present approaches represent ‘genetic marker’ approaches. These serve to employ linkage disequilibrium measures to use variants at a defined genomic position to infer unobserved variants at neighboring positions (Tewhey et al. 2012; Hoehe 2003).

The fraction of *cis* configurations (‘*cis* fraction’) (%) per individual genome is defined as the number of all autosomal genes with *cis* configurations divided by the total number of genes with ≥ 2 mutations, i.e. total configuration count; the *‘trans* fraction’ (%) per genome is complementary, 100% - *cis* (%). Thus, the *‘cis/trans* ratio’ per individual genome represents the ratio of *cis* fraction to *trans* fraction. The *‘cis/trans* ratio’ for a given population sample represents the median of the *cis* fractions (%) obtained across all genomes in relation to the median of their complementary *trans* fractions.

##### Scheme 2 Expected distribution of *cis*- and *trans* configurations of mutations in a population

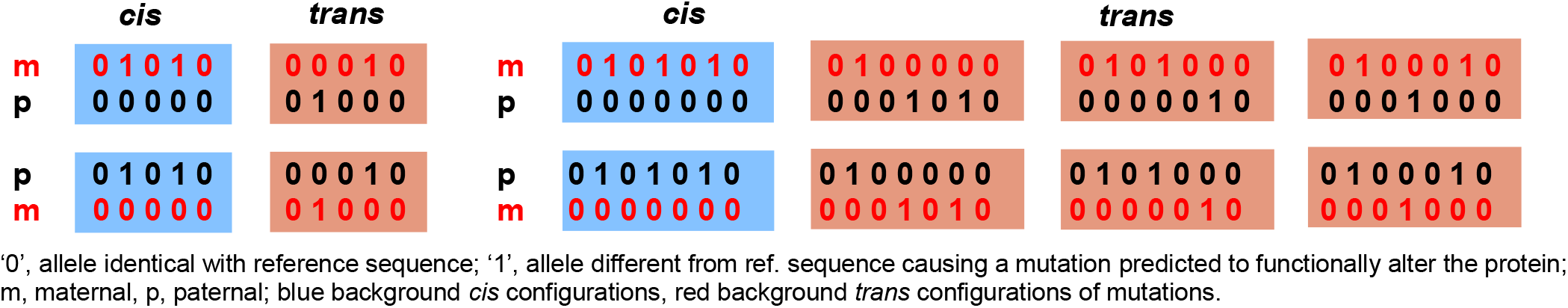

Scheme 2 provides an overview of all different *cis* and *trans* configurations expected to occur in a population, if the mutations (2 on the left, 3 on the right) in a protein-coding gene are distributed randomly between the maternal (m) and paternal (p) homologues. The number of different configurations for a defined number of n variants is 2^n^; as illustrated in Scheme 2, for n = 2 mutations, 4 different configurations would be expected, and for n=3 mutations, 8 configurations. Evidently, the same configuration type, i.e. distribution pattern, always occurs twice, with the parental origin interchanged, so these two should be biologically equivalent. Because current approaches to haplotyping do not allow distinction of maternal and paternal homologues, the maternal and paternal haplotypes are being collapsed into ‘Haplotype 1’ and ‘Haplotype 2’. Thus, in practice, we score a *cis* configuration, or any specific type of a *trans* configuration basically twice, leaving the relative fractions of *cis* and *trans* configurations constant. A *cis* configuration is scored, if ‘Haplotype 1’ has either solely ‘1’s or ‘0’s; if this is not the case, a *trans* configuration is scored. If the chance for every variant in a gene to occur on either homologue is equal, the expected fraction of *cis* configurations is calculated as 1/2, n being the number of variants. Obviously (see also Scheme 2), independently of the number of variants, there will always be 2 *cis* configurations, while the fraction of *trans* configurations grows exponentially (Graph).

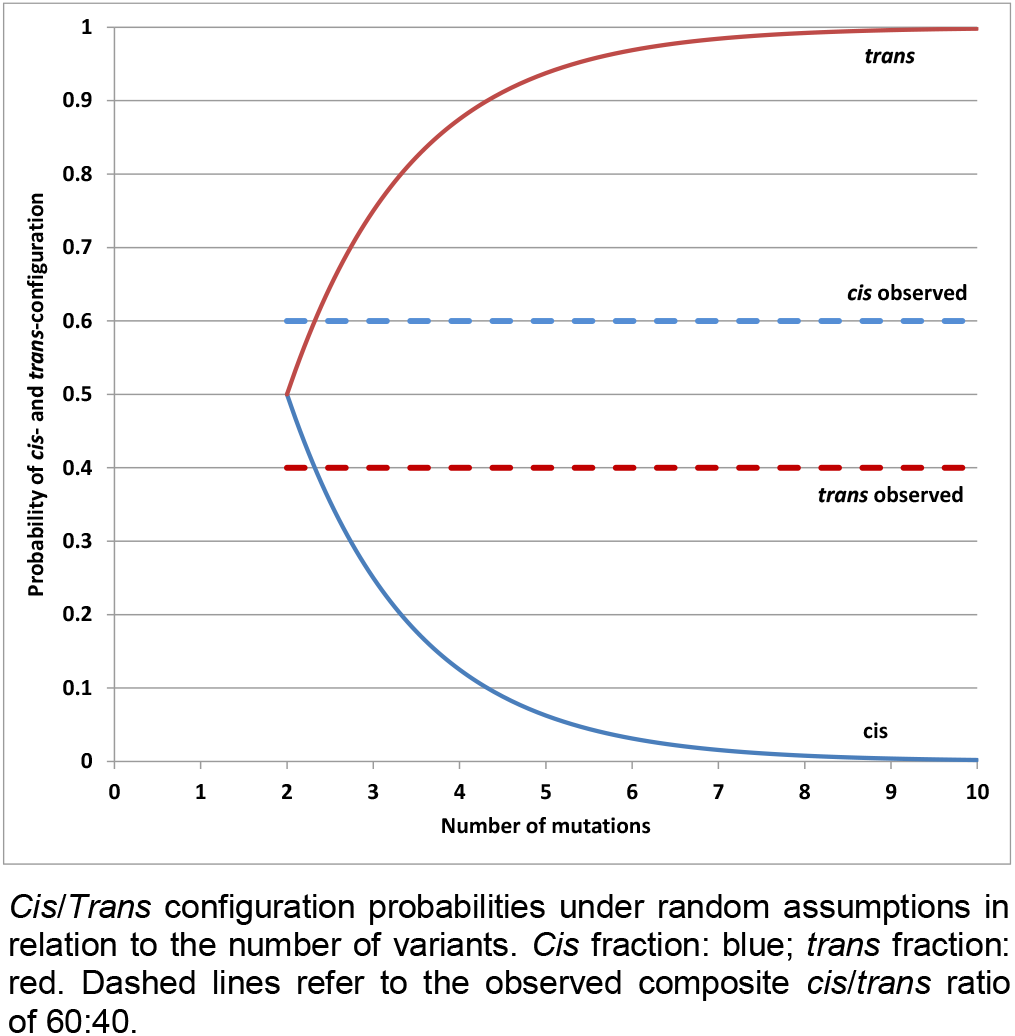

## Results

### Global abundance of *cis* configurations of mutations

*Cis* configurations of protein-altering mutations, leaving one form of the gene intact, would be expected to occur more frequently in human genomes to preserve organismal function (Hoehe et al. 2014). To test this hypothesis at large scale in 1,092 genomes (Abecasis et al. 2012) (Methods), we determined the ratio of *cis* to *trans* configurations of protein-altering mutations across all autosomal protein-coding genes for each of the genomes and derived the median of these ratios (Methods; see also Box 1). In fact, highly significant abundance of *cis* configurations of mutations was obtained, with a global *cis/trans* ratio of 59.6% to 40.4% (*P* < 3.53x10^−21^) (Table 1). Significant *cis*-abundance was moreover apparent in each of the four ancestry groups contained in the 1,092 genomes (Abecasis et al. 2012), with the *cis* fractions being similarly high in EUR (61.2%; *P* < 2.25x10^−21^), EAS (59.5%; *P* < 1.46X10^−17^) and AMR (60.1%; *P* < 3.09x10^−20^) and lower in AFR (54.7%; *P* < 1.66x10^−14^). An excess of *cis* configurations was also evident when examining the entirety of AA exchanges, with nearly identical results obtained from the total of 1,092 genomes *(cis/trans* ratio 59.7% to 40.3%, *P* < 6.16x10^−51^) as well as each of the four ancestry groups alone (Table 1). Furthermore, significant *cis*-abundance was observed for all sSNPs contained in the coding sequences, with an overall *cis/trans* ratio of 62.3% to 37.7% (*P* < 1.89x10^−71^); *cis* fractions were similar in EUR (62.9%, *P* < 1.55 x 10^−71^), EAS (62.9%, *P* < 4.56x10^−65^), and AMR (62.4%, *P* < 7.15x10^−71^), and lower in AFR (55%, *P* < 4.60x10^−43^) (Table 1). Notably, for calculation of significance values, we have taken into account that the probability to occur in *cis* by chance decreases with increasing numbers of variants in a gene (Methods; see also later chapter). *Cis*-abundance of protein-altering mutations was also evident when analyzing each of the 14 populations that were contained in the 1,092 genomes, with their *cis/trans* ratios being nearly identical with the median of their respective ancestry groups, with the exception of population ASW in AFR, its *cis* fraction of 56.3% slightly exceeding the AFR median (Supplemental Table S1). When analyses were expanded to include the entirety of AA exchanges and sSNPs, the results were once more highly congruent (Supplemental Table S1). Importantly, significant *cis*-abundance was cross-validated by the analysis of the 184 experimentally haplotype-resolved PGP genomes (Methods), with nearly identical ratios of 60.4% to 39.6% (*P* < 1.66x10^−16^) for protein-altering mutations (Methods), and ratios of 63% to 37% (*P* < 4.37x10^−51^) and 65.1% to 34.9% (*P* < 7.80x10^−70^), respectively, for the totals of AA exchanges and sSNPs (Table 1). In sum, these results show that protein-altering mutations, and coding variants as a whole, are in effect not distributed randomly between the two homologues of a gene, but do occur significantly more frequently on one of the two gene forms. On average, approximately 60% of the mutated autosomal genes in each genome have their coding variants in *cis*, and approximately 40% in *trans.*

**Table 1.** Global abundance of *cis* configurations of coding variants

| Population samples<sup>1</sup> | No. phased genomes<sup>1</sup> | *Cis* configs p-a mut<sup>2</sup> (%)<sup>3</sup> | *Trans* configs p-a mut<sup>2</sup> (%)<sup>4</sup> | *P*-value<sup>5</sup> | *Cis* configs AA exch<sup>6</sup> (%)<sup>3</sup> | *Trans* configs AA exch<sup>6</sup> (%)<sup>4</sup> | *P*-value<sup>5</sup> | *Cis* configs sSNPs<sup>7</sup> (%) | *Trans* configs sSNPs<sup>7</sup> (%) | *P*-value<sup>5</sup> |
| --- | --- | --- | --- | --- | --- | --- | --- | --- | --- | --- |
| Total | 1,092 | 59.6 | 40.4 | $3.53 \times 10^{-21}$ | 59.7 | 40.3 | $6.16 \times 10^{-51}$ | 62.3 | 37.7 | $1.89 \times 10^{-71}$ |
| EUR | 379 | 61.2 | 38.8 | $2.25 \times 10^{-21}$ | 60.7 | 39.3 | $4.84 \times 10^{-52}$ | 62.9 | 37.1 | $1.55 \times 10^{-71}$ |
| EAS | 286 | 59.5 | 40.5 | $1.46 \times 10^{-17}$ | 60.2 | 39.8 | $2.99 \times 10^{-45}$ | 62.9 | 37.1 | $4.56 \times 10^{-65}$ |
| AMR | 181 | 60.1 | 39.9 | $3.09 \times 10^{-20}$ | 59.5 | 40.5 | $3.34 \times 10^{-50}$ | 62.4 | 37.6 | $7.15 \times 10^{-71}$ |
| AFR | 246 | 54.7 | 45.3 | $1.66 \times 10^{-14}$ | 53.5 | 46.5 | $2.43 \times 10^{-30}$ | 55.0 | 45.0 | $4.60 \times 10^{-43}$ |
| PGP | 184 | 60.4 | 39.6 | $1.66 \times 10^{-16}$ | 63.0 | 37.0 | $4.37 \times 10^{-51}$ | 65.0 | 35.0 | $7.80 \times 10^{-70}$ |

<sup>1</sup> 1,092 statistically haplotype-resolved genomes from the 1000 Genomes (1000G) Consortium including the four different ancestry groups EUR, EAS, AMR, AFR described by Abecasis et al. (2012); 184 experimentally haplotype-resolved genomes from the Personal Genome Project (PGP) described by Mao et al. (2016).
<sup>2</sup> Mutations predicted to alter the protein, 'protein-altering mutations', from 1000G annotation database (Abecasis et al. 2012), or annotated using a combination of PolyPhen-2 (Adzhubei et al. 2010), SIFT (Ng and Henikoff 2001) and GERP conservation scores (Cooper et al. 2005) in the PGP genomes.
<sup>3</sup> Values represent the median of *cis* fractions (%), i.e. the number of *cis* configurations divided by total configuration count per genome across the genomes.
<sup>4</sup> Analogous to <sup>3</sup>, *trans* fractions (%) being 100% - *cis* fractions (%).
<sup>5</sup> Exact Binomial test (see Methods; Hoehe et al. 2014); *cis* counts were tested against the null hypothesis with a (composite) probability $P = 0.4$ for a *cis* configuration.
<sup>6</sup> Amino acid (AA) exchanges from 1000G annotation database (Abecasis et al. 2012), or the PGP database, respectively.
<sup>7</sup> Synonymous SNPs from 1000G or PGP database.
Configs, configurations; p-a, protein-altering; mut, mutations, AA exch, amino acid exchanges

### More *cis* than *trans* configurations in virtually all individual genomes

Inspecting the 1,092 genomes individually, in fact, 99.7% of the 1,092 genomes exhibited more *cis* than *trans* configurations of protein-altering mutations. Individual *cis* fractions varied within a limited range, between 55.7 and 67.1% in EUR and at most between 52.5 and 67.4% in AMR (Fig. 1A). Inspecting AFR, individual *cis* fractions were between 49.7 and 63.1%, with nearly half exceeding 55%. Testing each of the 1,092 genomes for statistical significance of *cis*-abundance using an exact binomial test (Methods), literally every single genome exhibited significantly higher *cis* fractions than would be expected if the mutations were distributed randomly between the two homologues of a gene (*P* < 2.55x10^−29^-2.30x10^−10^ in EUR, EAS and AMR genomes, 8.83x10^−23^-1.32x10^−07^ in AFR genomes). Examining the *cis* and *trans* configurations that were constituted by the entirety of AA exchanges, the results were highly similar (Fig. 1B): individual *cis* fractions ranged between 54.5 and 64.1% in EUR, EAS and AMR, and between 49.3 and 63.2% in AFR. *Cis*-abundance was again highly significant in each of the 1,092 genomes, with *P* < 1.04x10^−67^-2.38x10^−31^ in EUR, EAS and AMR genomes, and *P* < 4.76x10^−60^-3.68 x 10^−15^ in AFR genomes. Finally, also the *cis* configurations of coding sSNPs were significantly in excess in each of the 1,092 genomes, with fractions between 57.0 and 66.7% in AMR, EAS and EUR (*P* < 5.54x10^−94^-7.95x10^−53^) and between 52.1 and 63.1% in AFR (*P* < 9.18x10^−69^-7.32 x10^−27^) (not shown). The seemingly lower variance of *cis* and *trans* fractions between individual genomes in the case of AA exchanges and sSNPs was obviously due to the roughly 3-fold higher number of phase-sensitive genes per genome in these cases. Also, each of the 184 experimentally haplotype-resolved PGP genomes exhibited significantly higher *cis* than *trans* fractions for each type of coding variants. Specifically, the individual *cis* fractions of protein-altering mutations varied between 49.4% and 70.8% (*P* < 2.54x10^−26^-7.99x10^−06^) (Fig. 1C). Analyzing all AA exchanges, individual *cis* fractions were between 56.5% and 66.9% (*P* < 2.46x10^−61^-1.00x10^−38^) (Fig. 1D), and the highest in the case of sSNPs, between 57.0 and 69.2% (*P* < 3.87x10^−93^-9.76x10^−42^). In sum, these results strongly support the assumption that significant abundance of *cis* configurations of protein-altering mutations, and of any class of coding variants in general, represents a key characteristic of diploid human genomes. The observation that significant abundance of *cis* configurations is not restricted to protein-altering mutations suggests a mechanism other than purifying selection underlying the preferential location of mutations on the same chromosomal homologue of a gene. In this context it is important to note that significant *cis*-abundance was also obtained when analyzing all coding variants together, i.e. protein-altering mutations, other AA exchanges and sSNPs as they co-occur in many genes (*cis/trans* ratio 58.2% to 41.8%, *P* < 3.38x10^−89^). Supported by the decay of *cis* fractions in AFR, *cis*-abundance of coding variants most possibly reflects a yet unrecognized manifestation of linkage disequilibrium (LD) as a common, fundamental feature of genome variation.

**Figure 1.**
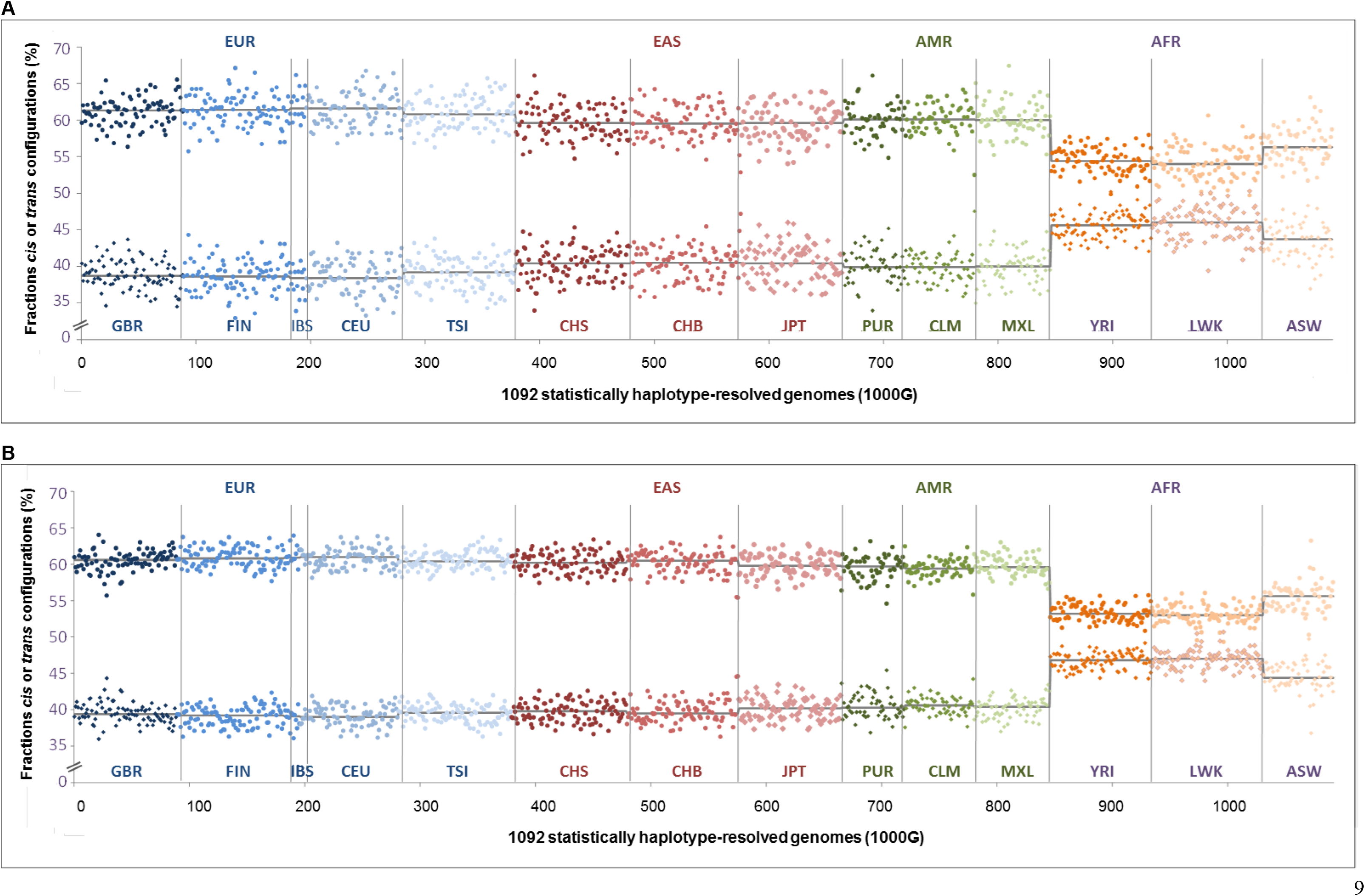
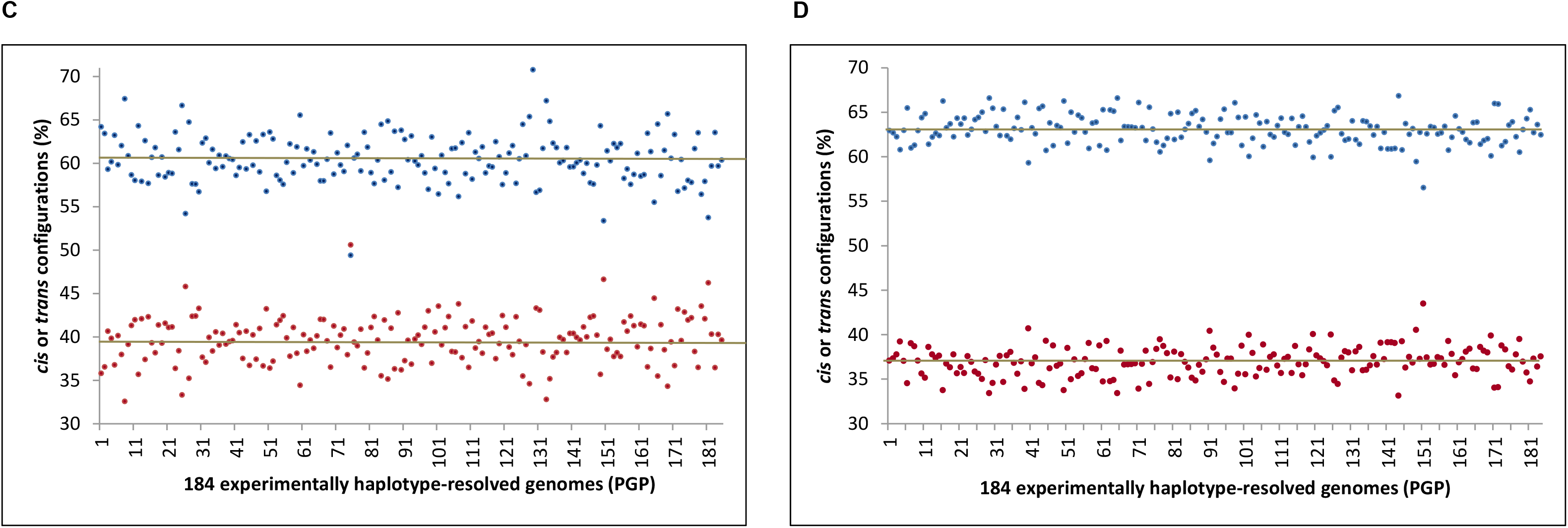
Individual fractions of *cis* and *trans* configurations of mutations in 1000 Genomes and PGP. (A) Individual fractions of *cis* and *trans* configurations of protein-altering mutations (%). In the upper half, the *cis* fractions are shown for each of the 1,092 statistically haplotype-resolved genomes from the 1000 Genomes (1000G) database (Abecasis et al. 2012), ordered by population (x-axis); individual *cis* fractions (%) (y-axis) represent number of *cis* configurations divided by total configuration count per genome. The complementary *trans* fractions (100%-*cis* fraction (%)) are shown in the lower half. Individual *cis* and *trans* fractions, respectively, are color-coded by ‘ancestry-based group’ as indicated on top, EUR (blue), EAS (red), AMR (green), and AFR (orange-brown). Within these groups, *cis* and *trans* fractions are further distinguished by population as indicated at the bottom, coded by main color nuances and separated by vertical lines. Median values (horizontal black lines) are shown for each of the fourteen populations; color-coding according to Abecasis et al. (2012). (B) Correspondingly, individual fractions of *cis* and *trans* configurations resulting from analysis of all AA exchanges. (C) Fractions of *cis* (blue) and *trans* (red) configurations of protein-altering mutations shown for each of the 184 experimentally haplotype-resolved PGP genomes. (D) Correspondingly, individual fractions of *cis* and *trans* configurations resulting from analysis of all AA exchanges. In contrast to (A) and (B), *cis* and *trans* fractions are color-coded.

### *Cis*-abundance primarily driven by pairs of mutations

The reported *cis/trans* ratios were obtained from genes with different numbers of variants. Because the probability of variants to occur in a *trans* configuration increases with the number of variants in a gene, these ratios essentially represent composite values. Therefore we dissected these further, assessing the *cis/trans* ratios separately for configurations with 2 up to 5 variants, which accounted for 95.7% of all configurations in the case of protein-altering mutations, and similar fractions in the case of AA exchanges and sSNPs, respectively. By far the most frequent configurations of protein-altering mutations, 66.7%, were found to consist of pairs of mutations, 65.8% of which resided in *cis.* The second most frequent configurations, 18.6%, were combinations of 3 mutations, occurring in equal proportions in *cis* or *trans.* As expected, the fractions of *trans* configurations increased to 53.4% in the case of 4 and 60.3% in the case of 5 mutations, accounting, however, for only small fractions of all configurations, 7.3% and 3.1%, respectively (Supplemental Table S2A). Essentially similar results were obtained when dissecting the *cis/trans* ratios determined from the entirety of AA exchanges and sSNPs; specifically, 57.7% and 59.6%, respectively, of all configurations consisted of pairs of variants, 67.9% and 69.9% of which resided in *cis.* Finally, analyzing all types of coding variants together, nearly half of all configurations consisted of pairs of variants, 68.7% of which were in *cis* (Supplemental Table S2B). Dissecting the composite *cis/trans* ratios separately in AFR, EUR, AMR and EAS showed an excess of pairs of protein-altering mutations also in each ancestry group: these were between 65.5% (AFR) and 67.7% of all configurations, of which 62.4 (AFR)-67.8% resided in *cis* (Supplemental Table S2A). Dissecting the composite *cis/trans* ratios in the PGP genomes, the pairs of protein-altering mutations accounted for nearly 65% of all configurations, of which 66.8% were in *cis* (Supplemental Table S2A). In sum, the dissection of the composite *cis/trans* ratios shows that the abundance of *cis* configurations is primarily driven by pairs of mutations that are predominantly in *cis.*

To estimate the excess of *cis* configurations we compared observed and theoretically expected *cis/trans* ratios. The probability *P* for a configuration with a defined number of *n* variants to reside in *cis* is 1/2^n-1^, if the chance for every variant in a gene to occur on either homologue is equal. So for pairs of coding variants, the observed *cis/trans* ratios were 66:34-70:30 versus expected 50:50; for combinations of 3 variants observed 50:50-54:46 versus expected 25:75, for combinations of 4 variants 45:55-47:53 versus 12.5:87.5, and for 5 variants 39:61-41:59 versus 6.25:93.75. This corresponded to a 1.32-1.4 up to 6.24-6.56-fold enrichment of *cis* fractions. Then we generated 1,092 virtual, phased genomes, assigning to each variant in a gene a 50:50 chance to exist on either homologue (Methods), and dissected the simulated data set accordingly. The simulated *cis/trans* ratios being virtually identical for 2 up to 5 variants to the expected ones (as calculated above), we derived the expected composite *cis/trans* ratios, ∼39:61 for protein-altering mutations, ∼37:63 for both AA exchanges and sSNPs, respectively, and 33:67 combining all types of coding variants (Supplemental Table S3). Thus, *cis/trans* ratios observed ∼60:40 versus expected below 40:60. These results had provided the basis for the use of an expected probability of 0.4 for a *cis* configuration to occur, when testing each of the 1,092 genomes for significance of the observed *cis* fraction. So even where *cis* ratios are minimally below 50%, which is the case in three of 1,092 genomes, and one of 184 PGP genomes, they are still significantly higher than would be expected by chance. These results underscore the overall tendency of coding variants to reside on the same homologue. Thus, diploid genes have two variable and potentially functionally altered homologues to a far lower extent than expected by chance.

### *Cis*-abundance driven by inter-mutation distance and frequency

In an attempt to explain observed *cis*-abundance, we considered ancestral admixture as a potential common underlying mechanism. Accordingly, pairs of protein-altering mutations in *cis* would be closely spaced and common, and pairs of mutations in *trans* distant and less frequent. So we compared, in a first step, the inter-mutation genome distances between totals of mutation pairs in *cis*, and *trans;* these are reported separately for EUR and AFR due to differences in the decay of LD. In effect, pairs of protein-altering mutations in *cis* were much closer together than pairs in *trans*, with inter-mutation distances in EUR of 1570 bp (median) versus 5290 bp, and 1830 versus 4771 bp in AFR. The same picture was observed when examining the distances between all pairs of AA exchanges and coding sSNPs, respectively: the variants in *cis* were much more closely spaced than the variants in *trans*, and the latter were less distant in AFR than in EUR; the absolute distances were, however, generally larger (Supplemental Table S4). Overall, pairs of variants in *cis* extended over genome distances up to nearly 3,900 bp (median), and in *trans* up to nearly 8,000 bp in EUR and ∼7,200 bp in AFR (Supplemental Table S4). These results were confirmed by analysis of PGP genomes, with inter-mutation genome distances of 2,584 bp (median) between pairs of protein-altering mutations in *cis*, and of 5,984 bp between pairs in *trans.* Examining the distances between all pairs of AA exchanges and sSNPs, respectively, the variants residing in *cis* spanned genome distances up to 3,562 bp, and the variants in *trans* distances up to 8,280 bp, largely consistent with the intervals estimated in 1000G (Supplemental Table S4). Where pairs of protein-altering mutations in *cis* and pairs of coding sSNPs in *cis* were found to co-exist within a diploid gene, which was the case for roughly 12-15 genes per genome, they were found to reside on the same homologue in over 83% of cases, indicating localization within the same ancestral segment.

In a second step, we examined the relationship of inter-mutation distance with *cis/trans* ratio. *Cis* and *trans* configurations were sorted by inter-mutation distance, binned per 6,000 configurations, and an average inter-mutation distance with its corresponding *cis/trans* ratio assessed per bin. Accordingly, the smallest distance in EUR, 11 bp, corresponded to the highest *cis/trans* ratio, 77:23, which declined to 51:49 at a distance of 54,281 bp and fell marginally below 50% at the largest average inter-mutation distance calculated, 81,099 bp, where a cumulative *cis* fraction of 67.8% was reached (Supplemental Fig. S1A, B). A similar inverse relationship between average inter-mutation distance and *cis/trans* ratio was observed in AFR, where an increase of distances from 14 to maximally 57,466 bp correlated with a (more rapid) decrease of *cis/trans* ratios from 75:25 to 48:52, with a cumulative *cis* fraction of 62.4% (Supplemental Fig. S1C, D). Analogous analyses performed with the entirety of AA exchanges and coding sSNPs (Supplemental Fig. S1E-H) strongly confirmed this picture. Taken together, significant *cis*-abundance is primarily due to an excess of pairs of mutations in *cis* that are closely spaced, while pairs of mutations farther apart are more likely to reside in *trans.* This suggests that *cis*-abundance is largely driven by distance.

In a third step, we compared the frequencies of pairs of protein-altering mutations in *cis* versus *trans.* To this end, we determined first the different, unique pairs of mutations that existed in either configuration, for each of the phase-sensitive genes in both EUR and AFR. Then we calculated for each pair the average mutation frequency, averaging the minor allele frequencies (MAFs) of each of the two mutations which were provided by the 1000G database for each ancestry group. Notably, the mutations occurring in pairs were found to exhibit very similar frequencies, and combinations of common mutations with singleton mutations, for instance, were very rarely observed. To begin with, we compared the average mutation frequencies between the totals of mutation pairs in *cis* versus *trans.* These (*quasi* the averages across the average mutation frequencies) were 0.18 and 0.16, respectively, in EUR, and 0.143 and 0.128 in AFR. Then we comparatively evaluated the average mutation frequency spectra. Overall, the frequencies of the mutation pairs in either configuration in both ancestry groups peaked between 0.1 and 0.25 (Supplemental Fig. S2A, B), in striking contrast to the MAF spectrum obtained from the entirety of protein-altering mutations in each of the ancestry groups, where 56% of all mutations in EUR (45% in AFR) had an MAF ≤ 0.01 (Supplemental Fig. S2C). Then, the average mutation frequency spectrum of pairs in *cis* in EUR peaked between > 0.2 and ≤ 0.25, and pairs in *trans* between > 0.15 and ≤ 0.2 (Supplemental Fig. S2A). In AFR, both frequency spectra were highest between > 0.1 and ≤ 0.15 (with pairs in *trans* exceeding *cis*), mutation pairs in *cis*, however, were twice as frequent as pairs in *trans* between > 0.2 and ≤ 0.25 (Supplemental Fig. S2B). So pairs of protein-altering mutations in *cis* overall are more frequent than pairs in *trans.* In sum, these findings support ancestral admixture as a potential explanation for observed, significant *cis*-abundance. Accordingly, this phenomenon results primarily from pairs of protein-altering mutations, and coding variants as a whole, that are closely spaced and therefore have been inherited together until present. Thus, they are more common than pairs of mutations in *trans* which, much farther apart, have been subject to recombination, and also may be due to evolutionary forces other than recombination, such as genetic mutation and positive selection. So pairs of co-occurring protein-altering mutations in *cis*, which are even closer together than other pairs of coding variants in *cis*, may represent ancestral signals of potential functional significance, and mark small ancestry segments in the ‘mosaic that is the human genome’ (Pääbo 2003).

Rather than proceeding with in depth analyses to further elucidate the admixture processes generating *cis*-abundance of coding variants, we have chosen to prioritize analyses that address the potential biological importance of this phenomenon. In other words, rather than addressing aspects of the history of our species, we have chosen to address primarily the result of this history as it may shape gene and genome function. So we focus in the second part of analyses predominantly on the genes that have mutations predicted to alter protein function in either *cis* or *trans* configurations (where phase may have an impact) and underlie the observed *cis/trans* ratio of ∼60:40. Importantly, the internal consistency of described results and the fact that we have obtained nearly identical *cis/trans* ratios from entirely different wet lab and bioinformatics pipelines earlier (Hoehe et al. 2014) apparently validates the annotation of mutations used here, provided by the 1000 Genomes database or the application of PolyPhen-2 (Adzhubei et al. 2010) and SIFT (Ng and Henikoff 2001) as well as GERP conservation scores (Cooper et al. 2005) in the case of PGP (Methods).

### A global set of phase-sensitive genes

First of all, how many, and which genes have two or more protein-altering mutations, which could exist in either *cis* or *trans* configurations and constitute observed *cis*-abundance? Each of the 1,092 genomes had between 393 and 710 such phase-sensitive genes (median 487), equivalent to 2.2-3.9% of all autosomal protein-coding genes (Hg19, RefSeq). In 221-397 (median 297) of these genes, the mutations resided in *cis* and in 132-342 (median 193) in *trans;* for corresponding results for each of the four ancestry groups see Supplemental Table S5A, for corresponding numbers from all AA exchanges, which were overall ∼2.7-fold higher and affected 5.7-10.3% of all autosomal genes, see Supplemental Table S5B. Taken together, the relatively low level of variation in the fractions of *cis* and *trans* configurations observed across the genomes corresponds to a relatively limited range of variation in the numbers of phase-sensitive genes, and their *cis* and *trans* forms. These numbers were directly proportional to the numbers of protein-altering mutations in a genome (median 2869) across all ancestry groups. A detailed description of the entire numerical framework that defines the highly specific relationships between mutations and the different categories of mutated genes, as well as the simulation of the proportionality constants through random distribution of mutations onto existing exome structure, is provided in Supplemental Table S6A-D; Supplemental Table S7; see also Supplemental Note. In sum, at the bottom of observed *cis*-abundance are highly similar numbers of mutations per genome which generate highly similar numbers of phase-sensitive genes, and their *cis* and *trans* forms. Thus, to which extent are the genes in each of these categories the same? In other words, is there a common, shared set of phase-sensitive genes? Can we further distinguish within such a set two groups of genes, which have preferentially *cis*, or *trans* configurations?

In fact, we were able to identify a set of 2,402 genes with protein-altering mutations in either *cis* or *trans* configurations, which were shared by all ancestry groups (Fig. 2; Supplemental Table S8). To this end, we intersected the genes which were phase-sensitive in each of the ancestry groups, 4,000 in EUR, 3,357 in EAS, 4,005 in AMR and 5,217 in AFR, any two of which overlapped by 80-87%. This ‘global set of phase-sensitive genes’ unveiled a highly significant overrepresentation of manifold gene ontology (GO) terms (Methods) including receptor activity, mainly olfactory, *trans*-membrane signaling and G-protein coupled receptor (GPCR) activity, the detection of (chemical) stimuli and sensory perception, membrane-and extracellular matrix (ECM)-related components/molecules, cell adhesion, the binding of odorants and antigens, multiple activities involved in the metabolism of substrates and xenobiotics, and MHC proteins (*P* < 6.12x10^−46^-9.56x10^−04^ for the top 99 GO groups) (Supplemental Table S9A). These results were complemented by a highly significant enrichment of pathways (Methods) including the transduction of olfactory signals, GPCR signaling and other types of signal transduction, ECM organization/receptor interactions, cell surface interactions, numerous immune-related processes such as antigen processing, Graft-versus-host and autoimmune diseases as well as multiple other diseases, chemical carcinogenesis, numerous biosynthetic and metabolic pathways, e.g. xenobiotics, drug metabolism and transport (*P* < 1.31x10^−52^-9.4x10^−03^ for the top 61 pathways; Supplemental Table S9B). 68% of the genes contained in the global set were also found to be phase-sensitive in the 184 experimentally haplotype-resolved genomes. This subset of 1,627 experimentally haplotype-resolved genes was found significantly enriched for similar biological functions. PGP genes shared 78% of the overrepresented GO terms (*P*<1.30x10^−41^-0.001) and 76% of the pathways (*P*<6.03x10^−46^-0.01) with the global set (Supplemental Table S9C, D), allowing extraction of essentially the same biological results (Supplemental Fig. S3A, B). Taken together, there exists a common, global set of phase-sensitive genes encoding two different, potentially functionally distinct homologues. These may modulate cell-cell communication, cell-environment interactions and membrane-related processes, immune processes, metabolism and biosynthesis, and the development of disease.

**Figure 2.**
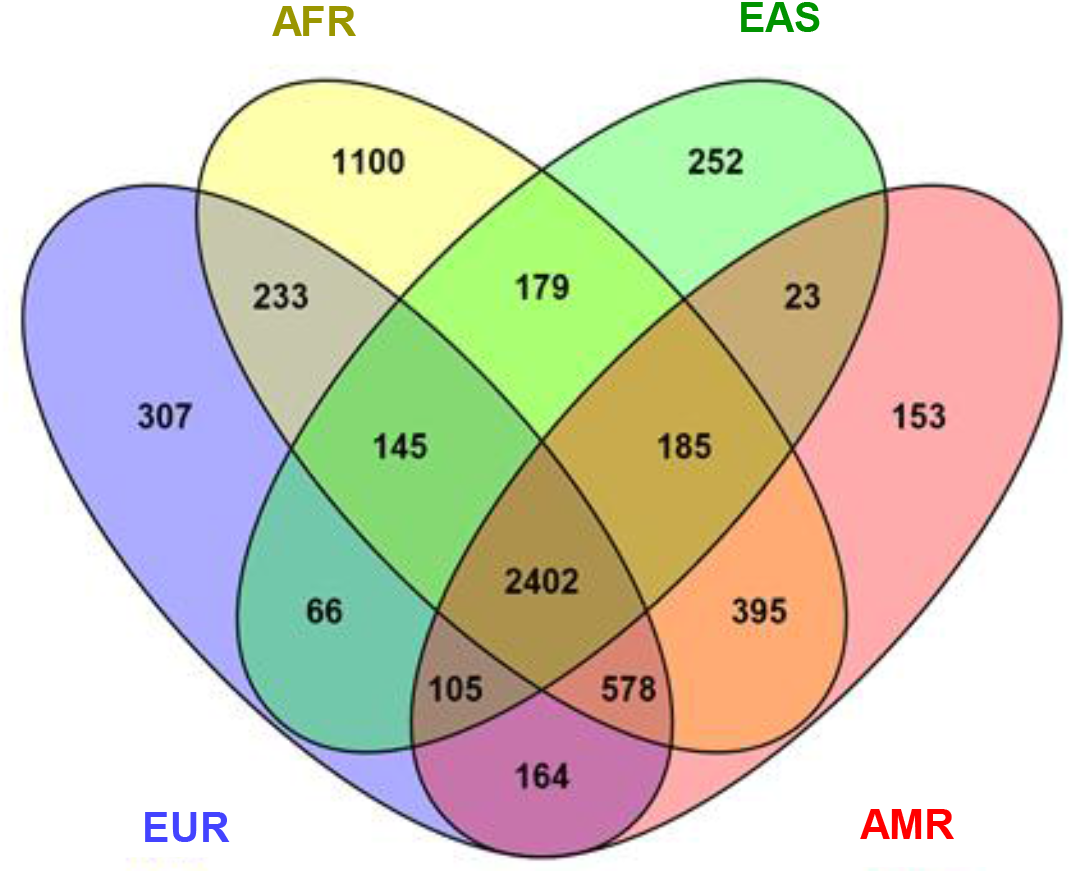
A global set of phase-sensitive genes. Venn diagram showing the set of 2,402 phase-sensitive genes (i.e. genes that have . 2 protein-altering mutations), which is shared by all four ancestry groups AFR, EAS, AMR and EUR (as color-coded) from the 1000 Genomes database. In addition, the numbers of phase-sensitive genes that are common to any 3 or 2 of these ancestry groups are indicated as well as those that are private to one of these ancestry groups.

In this work, we have focused on genes with ≥ 2 mutations to examine the distribution of mutations between the two homologues and global patterns of phase in the diploid human genome. Further analyses unveiled that the global set of phase-sensitive genes is an integral, though substantial part of a larger global set of genes with ≥ 1 protein-altering mutations (Supplemental Table S8) which similarly showed an overrepresentation of described functional categories (Supplemental Table S10A, B). The disproportionally larger contribution of the genes with ≥ 2 mutations compared to genes with 1 mutation, respectively, to the pathways and GO terms which were significantly enriched, are shown in Supplemental Fig. S4A, B (see Supplemental Note for further detail). Thus, the biological functions under consideration appear to be preferentially controlled by variable genes *per se*, ‘molecular diplotypes’ (Hoehe et al. 2014). In contrast, a set of 5,040 genes was identified (Supplemental Table S8), which did not contain any protein-altering mutations at all in the 1,092 genomes and was significantly enriched for numerous essential cellular functions which may not tolerate variability, including pathways involved in cell cycle, gene regulation and epigenetic mechanisms (*P* < 6.15x10^−25^-9.51x10^−11^ for the top 50 pathways; *P* < 2.35x10^−24^-2.12x10^−14^ for the top 50 GOs) (Supplemental Table S11A, B). Thus, the protein-altering mutations giving rise to both the global set of phase-sensitive and global set of variable genes are not randomly distributed across the exome, as in principle also shown earlier (Hoehe et al. 2014). Thus, evolutionary processes such as balancing selection may have resulted in the maintenance of long term diversity of these genes, preserving their functional flexibility to allow adaptation of the organism to internal and external stimuli and changes of environment. This assumption is supported by significant enrichment of the global set of phase-sensitive genes and, to a lesser extent, the expanded set of variable genes, with genes reported to be under balancing selection (BS) (*P* = 9.29x10^−07^ and *P* = 4.08x10^−03^, respectively) (Savova et al. 2016). Moreover, both global sets were found significantly enriched for monoallelically expressed (MAE) genes compared to biallelically expressed genes (BAE) (Methods; *P* = 1.75x10^−11^ and 1.99x10^−14^, respectively) (Savova et al. 2016), further corroborating their potential role as major modulators and adaptive agents. A potential evolutionary significance of these gene sets is furthermore indicated by significant enrichment with genes with ancient derived protein-coding polymorphisms or haplotypes predating the human-Neanderthal split (HNS) (Methods; *P* = 3.09x10^−16^ and *P* = 4.77x10^−12^, respectively) (Savova et al. 2016). The significance of enrichment with genes indicating evolutionary significance (BS, HNS and genes with evidence of human-chimpanzee *trans*-species polymorphisms (TSP)) is further enhanced when analyzing the set of 1,627 cross-validated phase-sensitive genes which is shared by PGP and the global set of phase-sensitive genes (*P* = 1.23x10^−09^ for BS, *P* = 6.56 x 10^−22^ for TSP, and *P* = 3.79 x 10^−22^ for HNS; see also Supplemental Fig. S5). Where the genes have ≥ 2 mutations the question arises, how are these mutations distributed between the two homologues of the genes in a population? Can we further distinguish within identified global set of phase-sensitive genes two groups of genes, which have either preferentially *cis*, or *trans* configurations of protein-altering mutations? And could these influence the biological functions in question differentially?

### *Cis*- versus *trans*-abundant genes

To this end, we evaluated first each of the 2,402 phase-sensitive genes across the entirety of 1,092 genomes using a Binomial test (Methods). In effect, we identified a subset of 1,227 genes exhibiting significantly more frequently *cis* configurations (1.05x10^−63^-0.048), which we defined as ‘cis-abundant genes’, and 786 genes having significantly more frequently *trans* configurations (1.78x10^−15^-0.049) defined as ‘*trans*-abundant genes’ (Supplemental Table S12A, B). The remainder, 385 genes (16%), had nearly equal proportions of both configurations. In a second approach, we determined first the significantly *cis*- and *trans*-abundant genes separately for each ancestry group, which we intersected to derive global sets of *cis*- and *trans*-abundant genes. These were identical with the gene sets derived in the first pass, confirming that the specific ‘configuration type’ of a gene is the same in all population samples. Thus, *cis*- or *trans*-abundance could represent a constant characteristic of a gene. In sum, the vast majority (84%) of the autosomal genes with ≥ 2 protein-altering mutations that were contained in the global set can be classified into the two major categories *cis*- and *trans*-abundant genes. These may reflect different mechanisms to exert and regulate gene functions.

So we examined, whether these *cis*- and *trans*-abundant genes could be differentially involved in the gene functions (GO terms) and pathways found overrepresented in the global set. In effect, they were found distinctively enriched for a number of pathways: *cis*-abundant genes for signaling by GPCR and signal transduction, biosynthetic and especially metabolic pathways, e.g. xenobiotics metabolism including cytochrome P450-mediated oxidation and other phase 1-related processes (*P*<1.81x10^−51^-0.008); *trans*-abundant genes for numerous immune response-related processes and diseases as well as autoimmune diseases, viral and infectious diseases, cell surface interactions, ECM-receptor interaction, signaling pathways, and drug transport (*P*<3.26x10^−7^-0.009) (Supplemental Table S13A, B). Thus, the disease-related phenotypes found enriched in the global set appear to be primarily mediated by *trans*-configurations. Both *cis*- and *trans*-abundant genes were significantly over-represented in pathways involving the transduction of olfactory signals, GPCR downstream signaling, and ECM organization (Supplemental Fig. S6A). In sum, over 75% of the pathways found enriched in the global set of phase-sensitive genes could be further distinguished by their preferential involvement of *cis*- or *trans*-abundant genes. A similar picture was observed for GO terms: an overrepresentation of significantly *cis*-abundant genes was for instance observed for GPCR activity and signaling, odorant binding, and activities involved in the metabolism of substrates. *Trans*-abundant genes e.g. were enriched for MHC proteins/receptors, antigen binding, ECM structural constituent and organization, and cell adhesion (Supplemental Table S13C, D). Both *cis*- and *trans*-abundant genes were significantly enriched for instance in olfactory and *trans*-membrane (signaling) receptor activity, the detection of (chemical) stimulus, sensory perception and membrane-related components (Supplemental Fig. S6B). In sum, 57-83% of the GO terms, evaluated separately for their taxonomies, could be further differentiated with regard to their preferential involvement of either gene category. Thus, *cis* and *trans*-abundant genes may differentially influence pathways and gene functions.

This classification of autosomal genes has been derived from the global set of phase-sensitive genes, which effectively represents 46-72% of these genes within each ancestry group. To test whether this classification is also valid in a population as a whole, we examined each of these ancestry groups separately. Setting a threshold of 10 configuration counts per gene, 1,173 significantly *cis*- and 670 *trans*-abundant genes were obtained in EUR, 966 and 590 in EAS, 981 and 497 in AMR and 1,265 and 817 in AFR, accounting for 78-88% of all autosomal phase-sensitive genes. Essentially similar results were obtained using a threshold of 5% of total genome count. Subsequent analysis of the 184 experimentally haplotype-resolved PGP genomes also validated this classification; correspondingly, 83.5% of the phase-sensitive genes were grouped into 778 significantly *cis*- and 436 *trans*-abundant genes. In sum, these results underscore the existence of *cis*- and *trans*-abundant genes as major categories of variable autosomal genes. Evidently, the group of *cis*-abundant genes is always larger than the group of *trans*-abundant genes, with ratios between 1.55:1 and 2:1. Thus, significant *cis*-abundance is the net result of these two groups. Both *cis*- and *trans*-abundant genes were found distributed across the autosomes in varying mixtures, leading to autosomal *cis/trans* ratios between 53:47 (chr. 10) and 69:31 (chr. 14) (Supplemental Table S14A, B; Supplemental Note). Autosomes with disproportionately higher fractions of *trans*-abundant genes include in addition to chr. 10 the chromosomes 6 and 8, autosomes with a higher density of *cis*-abundant genes in addition chr. 20 and chr. 22. An overview of the ‘phase-sensitive genome’ is provided in Supplemental Fig. S7A, B.

Subsequently we examined whether the *cis*- and *trans*-abundant genes identified in the global set are verifiable in the experimentally haplotype-resolved genomes. Intersecting the 1,227 *cis*-abundant genes which were common to all ancestry groups contained in 1000G, with the 778 genes *cis*-abundant in PGP resulted in an overlap of 322 genes. Correspondingly, 153 of the 786 *trans*-abundant genes were also observed in PGP, a sample only one-sixth part of the size. Fig. 3 shows examples of these cross-validated genes, demonstrating that statistically and experimentally haplotype-resolved genomes essentially deliver the same results and, moreover, illustrating the key point that the majority of autosomal genes with ≥ 2 mutations have either *cis* or *trans* configurations in significant excess. Thus, notable fractions of *cis*- and *trans*-abundant genes exhibited solely *cis* or *trans* configurations, respectively, i.e. gene-based *cis* or *trans* fractions of 100%, such as *PON2, OR8D2*, or *MT1A* (Fig. 3). On average, gene-based *cis* fractions were 82% (1000G) and 90% (PGP) of total configuration counts of 23 and 24%, respectively. Gene-based *trans* fractions were 80 and 86% of total configuration counts of 19 and 22%, respectively; for a more detailed quantitative characterization of both gene categories see Supplemental Note. Moreover, an expanded analysis corroborated that *cis*- and *trans*-abundance in effect represents a constant characteristic in nearly 90% of the genes with only 8.7% of all *cis*- and 12.9% of all *trans*-abundant genes changing configuration type in PGP (see Supplemental Note). Examples of *cis*-abundant genes in Fig. 3 include members of pathways and GOs that were significantly overrepresented in this gene category, such as *CYP2D6, GSTZ1, MAP4* and *SULT1C3* involved in xenobiotics metabolism, pharmacokinetics, biosynthesis and metabolism in general, or *OR8D2* and *OR2M7* involved in olfactory transduction, receptor activity and signaling (by GPCR). Numerous *cis*-abundant genes have conserved protein domains, such as *ANKLE1, LRRK2, ADAM33, CCDC137*, or *ZNF546*, a C2H2 zinc finger (C2H2-ZNF) gene. Moreover, some *cis*-abundant genes represent members of gene families, which as a whole are *‘cis*-abundant’, that is, their members are significantly more often *cis*-abundant, such as the olfactory receptor (OR) gene family (*P* = 3.41x10^−05^), which overall contributes 0.87% to the global *cis* fraction of ∼60%. Examples include furthermore *cis*-abundant genes involved in common, complex and autosomal recessive diseases such as *CYP2D6, ADAM33, ADD1*, and *PON2*, and particularly prominent disease genes such as *BRCA1* and *LRRK2.* Examples for *trans*-abundant genes (Fig. 3) include genes involved in immune response, immune and autoimmune diseases, and viral and infectious diseases, such as *HLA-E, KIR3DL2, TPO* and *DDX58;* genes involved in ECM-organization and components such as *COL11A2*, in (transmembrane) signaling such as *P2RX7* and *CEP55*, and in cancer such as *MTA1* and *NAT2.* Some of these *trans*-abundant genes, too, are members of predominantly *trans*-abundant gene families, such as the histocompatibility complex (*P* = 2.49x10^−03^), and collagen family (*P* = 1.26x10^−02^).

**Figure 3.**
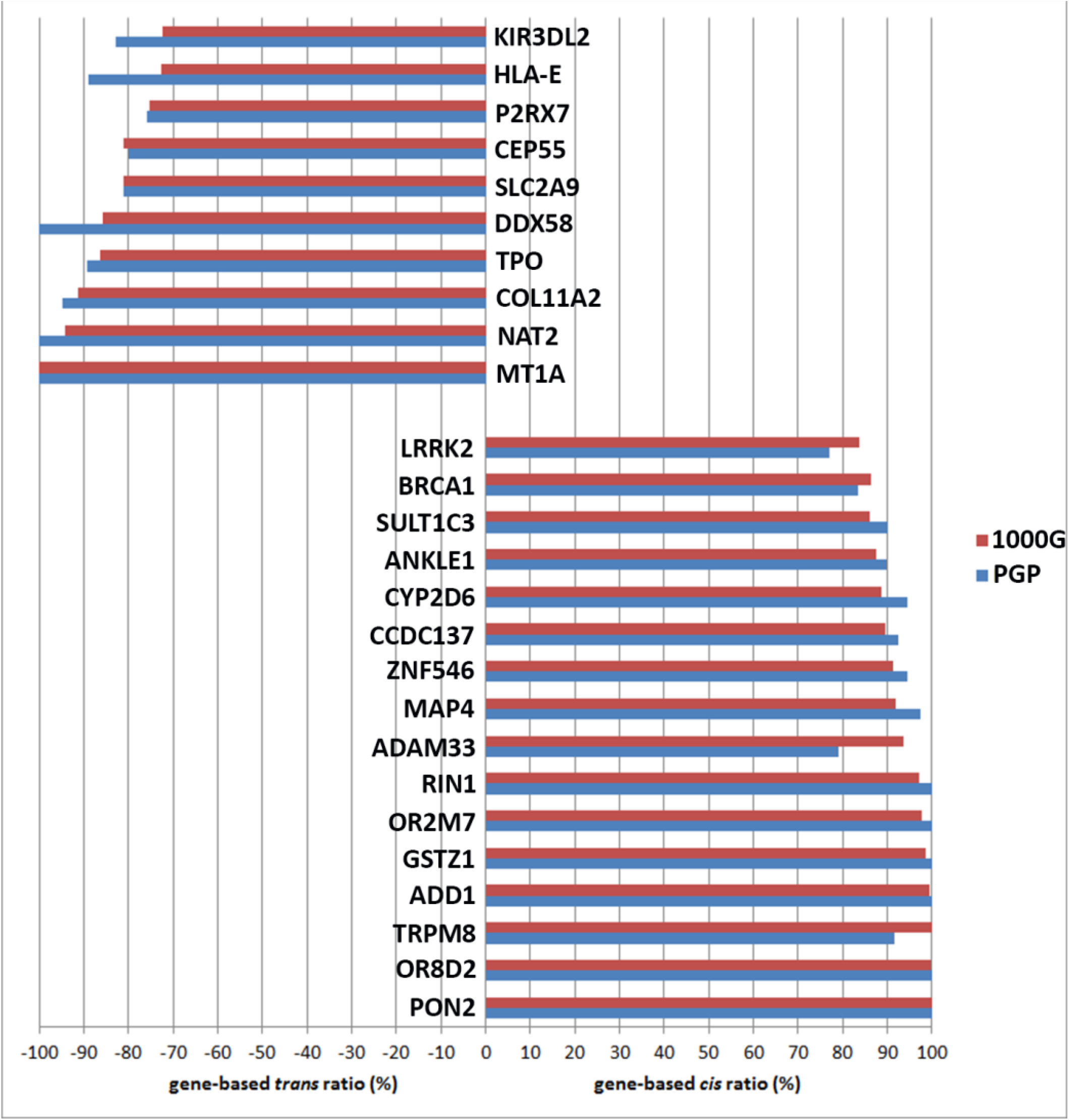
Examples of *cis*- and *trans*-abundant genes. Genes were selected from the sets of 322 *cis*- and 153 *trans*-abundant genes, which were shared by both 1000 Genomes (1000G) and PGP. Top left *trans*-abundant genes, lower right side *cis*-abundant genes; blue bars indicate the gene-based *cis*- and *trans* fractions (%) derived from the 184 experimentally haplotype-resolved PGP genomes, red bars the fractions derived from the 1092 statistically haplotype-resolved genomes from 1000G database. A gene-based *cis* ratio is defined as the number of *cis* configurations counted in a particular gene in a sample of genomes divided by the total configuration count of this gene in the sample; gene-based *trans* ratios are defined accordingly.

Subsequently we examined the specific positions of protein-altering mutations in *cis* versus *trans* across the genes. We attempted to identify different distributional patterns of the mutations, not only between, but also within the homologues, which could be functionally relevant. In a first step, we inspected the mutation genomic coordinates in several cross-validated genes in more detail and in fact observed striking differences. Thus, in *cis*-abundant genes such as for example *ZNF546* (Fig. 4A), or *ADAM33, PON2, ZNF626* and *KRT83*, there existed one pair of mutations that dominated the picture. Notably, the two mutations establishing such a ‘major configuration’ nearly always occurred together, and only rarely, one of the two protein-altering mutations occurred alone, or in combinations with other mutations. In addition to these major configurations, in case, small numbers of other pairs or combinations of mutations in *cis* or *trans* were observed (Fig. 4A). A much more mixed picture was observed in *trans*-abundant genes, as exemplified by *KRT3* (Fig. 4B). This was characterized by two or more, apparently less frequent pairs of mutations in *trans;* in this example, a *trans* pair existed to a small, but visible extent still in *cis.* To test whether these case observations represent a more general picture, we examined the 1,227 *cis*-abundant and 786 *trans*-abundant genes identified in the global set, as well as the 322 *cis* and 153 *trans*-abundant genes shared by both 1000G and PGP. We determined the proportion of the most frequent pair of mutations relative to the entirety of configurations in each of the genes, across the genomes. Then we binned the genes by percentage of their major configuration count. As demonstrated in Fig. 5A, a substantial fraction (31.2%) of the 1,227 *cis*-abundant genes exhibited one major pair of protein-altering mutations accounting for > 90 up to 100% of all *cis* configurations, and 42% one major pair accounting for > 80% of all *cis* configurations. This result was confirmed in the cross-validated genes (Fig. 5B, C). 12-20% of the genes had major pairs of mutations with frequencies of 20% or less; thus, these genes have mostly more frequent combinations of 3 and more mutations, such as *BRCA1.* A strikingly different picture was observed for *trans*-abundant genes, characterized by the absence of a major configuration which would be present in sizable fractions (Fig. 5D-F). In sum, significant *cis*-abundance essentially results from a considerable fraction of *cis*-abundant genes, which have highly frequent pairs of mutations co-occurring on the same homologue. Thus, we have traced *cis*-abundance back to specific, potentially ancient pairs of protein-altering mutations in *cis.* Attempting to interpret this finding, we considered epistatic interactions between these pairs of mutations, a potential role as compensatory mutations, or co-evolution. In order to explore whether these major, closely spaced mutation pairs could in fact indicate potential interactions between the mutations, we have mapped them onto protein tertiary structures using the Protein Data Bank (PDB) (Rose et al. 2013). Indeed, for some proteins, for example ESYT2 (Extended Synaptotagmin 2), a membrane-related protein, we found that the corresponding residue pairs existed in close physical distance (∼9 Å) (Fig. 6). This could suggest possibility of physical interaction and/or functional interdependence, for instance indicating coevolution (Anishchenko et al. 2017). Rounding up, we have obtained first, very preliminary evidence for potential functional implications of major mutation pairs in *cis*, the hallmark of *cis*-abundance.

**Figure 4.**
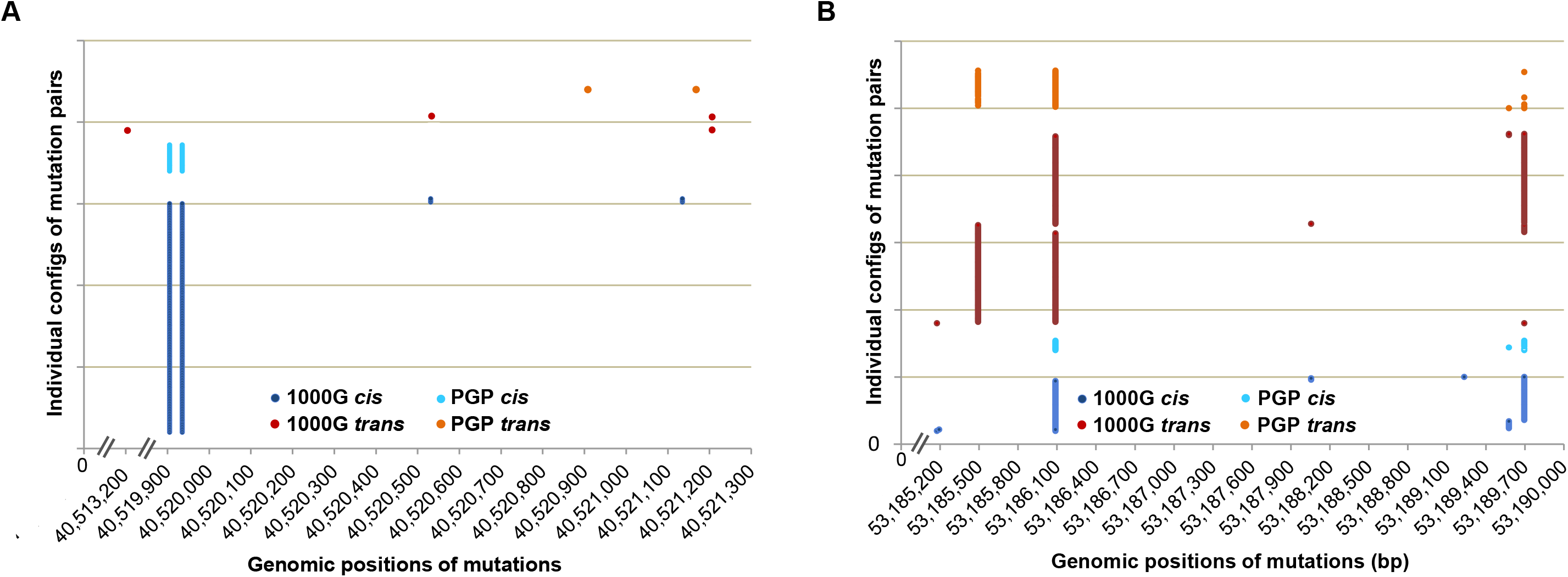
Typical configurations of pairs of protein-altering mutations in *cis*- and *trans*-abundant genes. The specific configurations of pairs of mutations, specified by genome positions (bp), are presented as observed in the 1,092 genomes from 1000G and 184 PGP genomes. Each individual pair of mutations is indicated by a pair of points, color-coded by source. (A) Distributional pattern of mutations in the *cis*-abundant *ZNF546* gene. The same pair of mutations in *cis* occurs in numerous individual genomes in both 1000G (blue) and PGP (light blue) and therefore appears as two horizontal, parallel lines; a different pair of mutations in *cis* is observed in 1000G in few individuals as well as two pairs of mutations in *trans*, which share the rightmost mutation; one pair of mutations in *trans* is observed in PGP. (B) Distributional pattern of mutations in the *trans*-abundant *KRT3* gene. Apparently, the pairs of mutations are less frequent; one pair of mutations occurs in *trans* in both 1000G and PGP (left); another pair of mutations (the left mutation shared with the other pair) occurs in *trans* in 1000G and in one individual in PGP; the same pair also shows a less frequent, though still sizable fraction of *cis* configurations in both 1000G and PGP, exemplifying the change from a *cis* to a *trans* configuration over increasing numbers of generations for more distant pairs of mutations.

**Figure 5.**
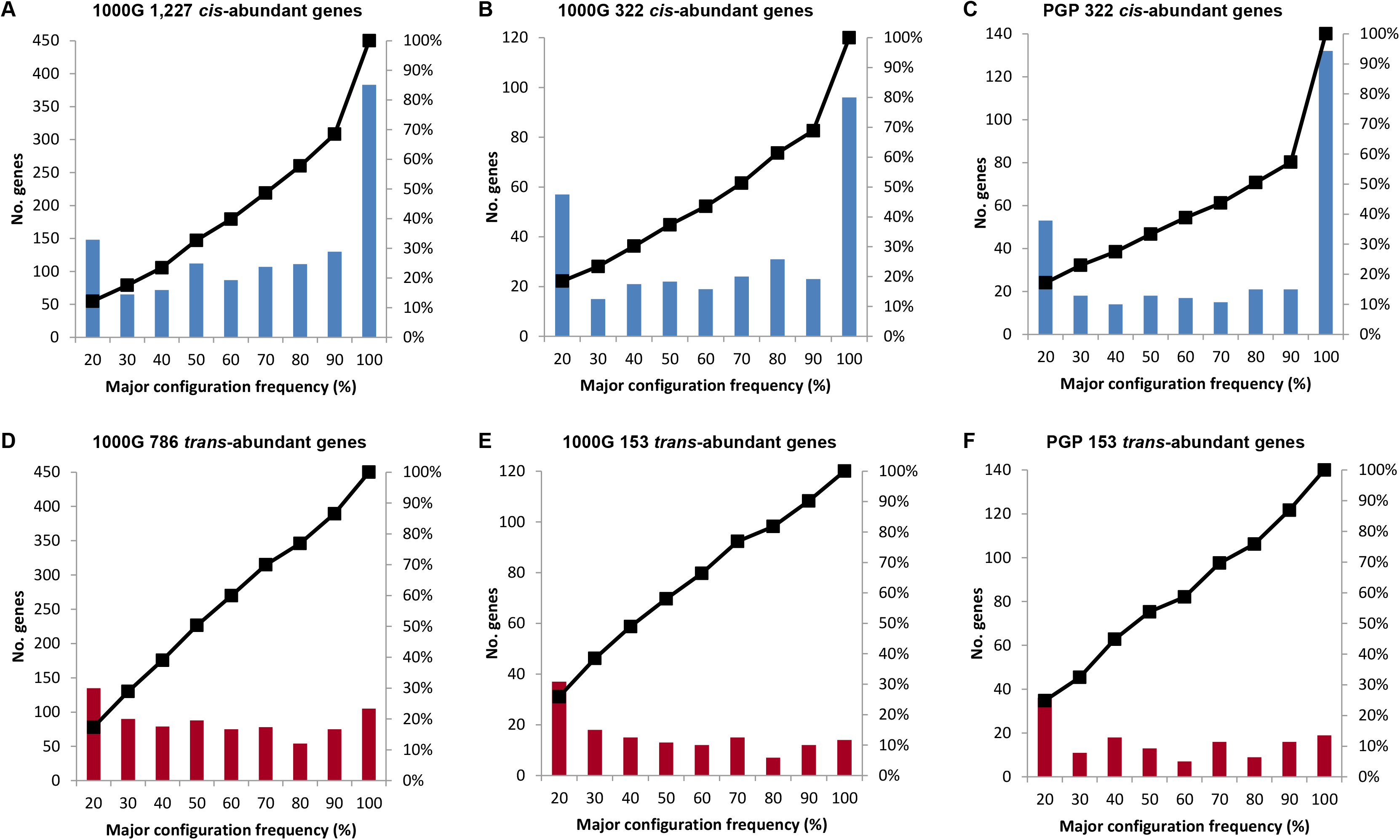
Major configuration frequencies in *cis*- and *trans*-abundant genes. The ‘major configuration’ of a *cis*- or *trans*-abundant gene is defined as the pair of mutations that exhibits the highest number of *cis, or trans* counts, respectively, relative to the total of *cis or trans* counts observed for this gene in a population sample. This ratio is defined as the ‘major configuration frequency’ (MCF) of a gene. The MCFs of defined numbers of genes are binned in 10% intervals. Specifically, they wer e assessed for 1,227 *cis*- and 786 *trans*-abundant genes in 1,092 genomes from the 1000 Genomes (1000G) database, and for the cross-validated sets of 322 *cis*- and 153 *trans*-abundant genes which were shared by both 1000G and PGP. (*A*) The blue bars indicate the numbers of *cis*-abundant genes (*y*-axis left) of a total of 1,227 *cis*-abundant genes, which have a major configuration with a frequency (%) as specified by given bins (*x*-axis). For example, referring to the highest blue bar on the right, 383 *cis*-abundant genes have a major configuration with a frequency between > 90 and 100% of total *cis* count. The black graph represents the cumulative percentage (%) of the number of these genes (*y*-axis right). (*B)* Accordingly, MCFs binned for 322 *cis*-abundant genes from 1000G and (*C)* for the identical 322 *cis*-abundant genes assessed in the 184 experimentally phased PGP genomes. The corresponding results for *trans*-abundant genes (red bars) are presented in (*D*), (*E*), and (*F*).

**Figure 6.**
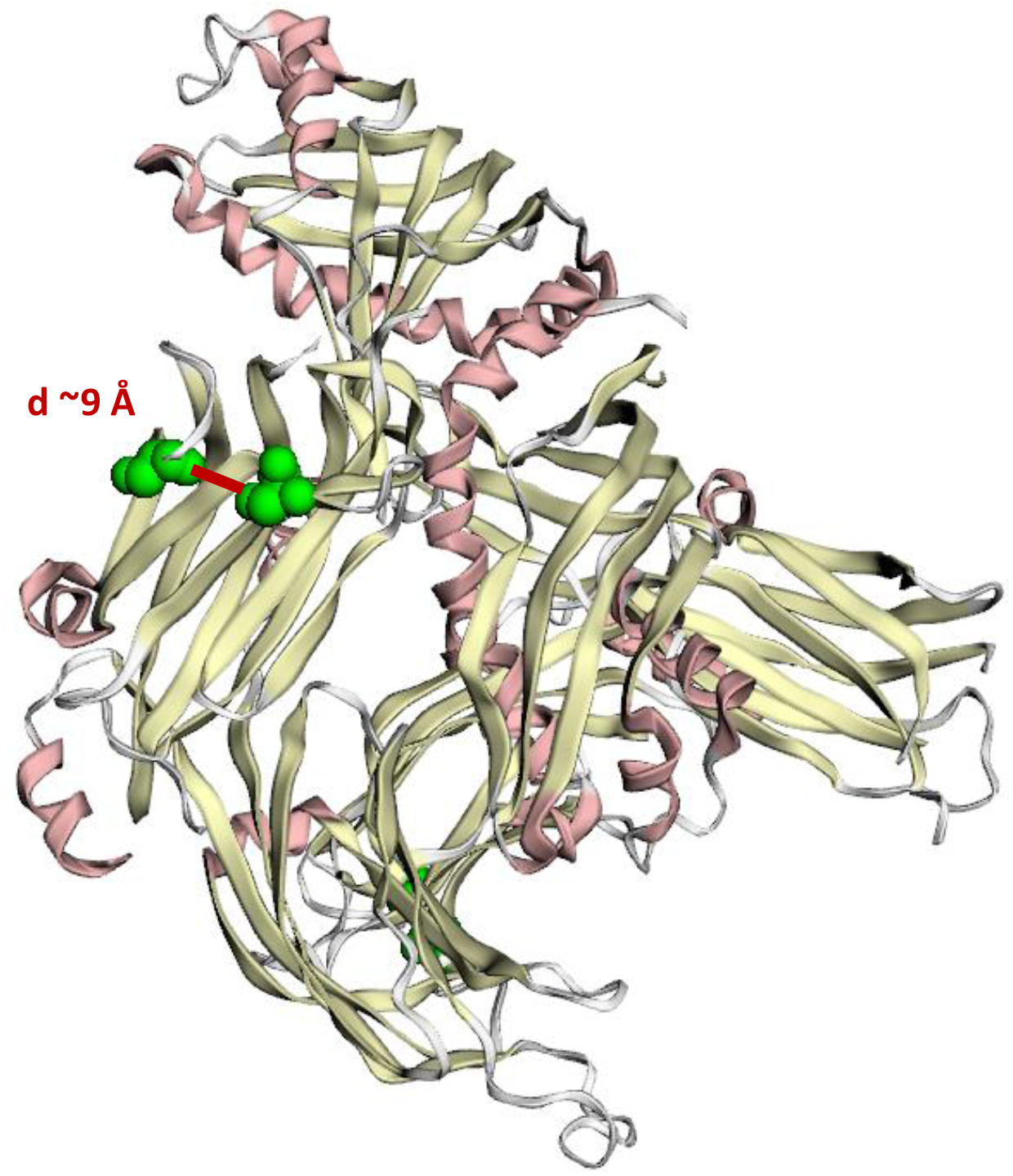
Major pair of mutations in a *cis*-abundant gene mapped onto protein tertiary structure. Two neighboring mutations (S584P and S610G) in the protein ESYT2 (Extended Synaptotagmin 2) are shown. Both mutations fall in the C2 domain that targets the protein to the cell membrane. Mutations are located within a physical distance of ∼9 Å, suggesting potential contact between the residue pair. Tertiary structure was taken from the PDB database (ID: 4P42). Mapping was generated with the MuPIT web server (Niknafs et al. 2013). Residue distance was computed with the Chimera software (Pettersen et al. 2004).

## Discussion

This work represents the first large scale analysis of both statistically and experimentally haplotype-resolved genomes, 1,276 in total from different populations, allowing establishment of common patterns of phase as key characteristics of diploid human genomes. Virtually all genomes showed significant abundance of *cis* configurations of protein-altering mutations, at an overall *cis/trans* ratio of about 60:40. Nearly all genes exhibited either *cis*, or *trans* configurations in significant excess, allowing distinction of *cis*- and *trans*-abundant genes as major categories of autosomal genes. These patterns were largely constituted by a common, global set of phase-sensitive genes that exist in either *cis* or *trans* configurations. The potential functional importance of these patterns was suggested by significant enrichment of this global set with gene sets indicating its involvement in adaptation and evolution; further by distinction of the functional classes concerned into those that are primarily mediated by *cis*-, or *trans*-abundant genes.

It is unlikely that these patterns represent an artifact due to bias in phasing. This statement is supported by (i) the high quality of 1000G statistical haplotype data and their strong validation by analysis of a substantial number of experimentally phased genomes; and (ii) the robustness of the experimental phasing method, ensured by an exceedingly low phasing error rate (Peters et al. 2012). Thus, additional steps to eliminate potential artifacts as causes of the *cis* and *trans* patterns of heterozygous variants, such as independent calculation of LD values from un-phased data, do not appear warranted. While some variation in the absolute numbers of *cis* and *trans* configurations of mutations has been observed, possibly due to differences in population structures and size, or annotation and validation procedures, the relative fractions of *cis* and *trans* configurations proved to be quite stable. Moreover, it appears unlikely that the *cis/trans* ratios of mutations predicted to alter the protein were biased by the annotation algorithms used (Hicks et al. 2011; Tennessen et al. 2012; Fu et al. 2013), as entirely different wet lab and bioinformatics pipelines, as well as control analyses of all AA exchanges and sSNPs produced highly similar ratios. Thus, the overall consistency of our data suggests that the key results can be considered robust and not an artifact of the specific algorithms and settings employed, or the parameters used.

This work represents an advance in the field of haplotype analysis in that an established experimental phasing method (Peters et al. 2012) has been applied at the population level to address ‘higher order questions’ concerning the diploid nature of the human genome (Wu and Dunlap 2002; Hoehe 2003; Suk et al. 2011; Tewhey et al. 2011). Recent efforts have mainly focused on refining and improving technologies for the efficient haplotyping of individual genomes. On the other hand, large scale efforts inferring haplotypes statistically from populations, like the HapMap and 1000 Genomes Project are primarily based on genetic marker concepts, i.e. using LD to infer the position of disease variants, while providing a catalogue of functional variants to help to prioritize candidate disease variants (Abecasis et al. 2012; Auton et al. 2015). In contrast, this work attempts to address the distributional patterns of mutations across genes, between and within the two homologous chromosomes and their potential functional consequences, to help to understand diploid gene and genome function. *cis*-abundance of protein-altering mutations and coding variants as a whole, represent yet another face of LD in the human genome, and the relationship between the extended haplotype blocks established by the HapMap or 1000 Genomes Project, and the phased exonic sequences remains to be determined. Essentially, our work shows how the patterns of LD in the human genome (Gabriel et al. 2002; Ardlie et al. 2002) affect the distribution of genetic variants between the two homologous chromosomes of autosomal genes. In this context, it is important to note that due to existing LD a ∼50:50 *cis/trans* ratio of the remaining classes of coding variants could not necessarily be expected in order to support functional significance of *cis*-abundance of protein-altering mutations. This issue is related to the challenge of distinguishing the causative variants from those in LD in the context of disease gene identification. Importantly, analogous analyses which we performed solely with AA exchanges other than those predicted to alter protein function, and sSNPs, failed to generate the results obtained with the protein-altering mutations.

Moreover, this work represents a first phase-informed analysis of protein-coding variation in the diploid human exome. Recent exome studies (Li et al. 2010; Tennessen et al. 2012; Fu et al. 2013; Lek et al. 2016; Telenti et al. 2016), in an effort to facilitate identification of potential disease-causing variants, have mainly focused on characterizing the functional spectrum of allelic variation in human genes and the underlying demographic and evolutionary forces that generated this variation. Issues of phase have yet remained unaddressed. Considering a total of 18,121 autosomal genes (RefSeq), phase may be of relevance for 6,284 genes, which have ≥ 2 protein-altering mutations in at least one of 1,092 genomes, while 6,797 genes have at most one mutation and 5,040 autosomal genes not any protein-altering mutations at all. Nearly half of all mutations predicted to functionally alter the protein in a genome (and over 60% of all AA exchanges) were found to exist in either *cis* or *trans* configurations and therefore require phase information. An overwhelming majority of all variable autosomal genes with ≥ 2 protein-altering mutations are either *cis*- or *trans*-abundant, and so are all genes with AA exchanges and sSNPs. Knowledge of phase is also indispensable in human exomes as there are many genes with mono-allelic or allele-specific expression, especially those modulated by epigenetic phenomena (Gimelbrant et al. 2007; Leung et al. 2015). In this case, the unique distribution of mutations across the two chromosomal homologues affects which mutations will ultimately have functional consequences. Thus, phase is an important link to differentially expressed transcriptomes and proteomes. Future questions to be addressed are which of the two homologues of *cis*-, and *trans*-abundant genes are expressed, in spatial and temporal context. The global set of phase-sensitive genes and with it, *cis*- and *trans*-abundant genes putatively encode two functionally distinct homologues, which particularly in combination with mono-allelic expression (MAE) have the potential to preserve functional flexibility, suggesting an important role of these genes for adaptation and evolution (see also Wu and Dunlap 2002; Chess 2012; Hoehe et al. 2014; Savova et al. 2016). These genes were found significantly enriched for functional categories such as cell-cell communication and cell-environment interactions, membrane-related processes, immune response, and metabolism and biosynthesis, supporting their importance for adaptation of organisms to changes in internal and external environment. Future studies will have to address the role of diplotypic genes in transcriptome, proteome, and phenotype diversity within and between cells, tissues, individuals, populations, species, at various developmental and physiological stages, as well as health and disease. Finally, the unprecedented nature of our cross-validated set of haplotype-resolved genomes provides a resource which allows linking each of the protein-coding homologues to transcriptional regulatory motifs and any non-coding elements that regulate their expression.

This work has introduced two major categories of variable autosomal genes: *cis*- and *trans*-abundant genes. The more frequent *cis*-abundant genes are characterized by common pairs of closely spaced protein-altering mutations. Thus, these may represent ‘evolutionary signals’ which trace back to ancient populations. Reasons for their preservation could be epistatic or compensatory interactions between these mutations, maintaining or enhancing the functionality of the protein (Kondrashov et al. 2002; DePristo et al. 2005; Ferrer-Costa et al. 2007; Baresic et al. 2010), co-evolution (Anishchenko et al. 2017), or hitchhiking effects (Slatkin 2008). Evidence for potential physical interaction between such mutation pairs in *cis* has been obtained in preliminary analyses at the example of ESYT2 (Fig. 6). Once the (available) 3D structures in protein databases are more complete, these seemingly old, co-occurring pairs of protein-altering mutations discovered in this work, may provide valuable information for the study of protein evolution and functionality (DePristo et al. 2005). *Trans*-abundant genes, on the other hand, apparently result from a mixture of mechanisms, such as the recombination of more distantly spaced mutations (exemplified by mutation pairs, which exist to a small percentage still in *cis*, while to a larger extent in *trans*), the occurrence of mutations, or positive selection such as in the case of HLA genes (Bustamante et al. 2005). This latter example referring to the relatively short HLA genes illustrates that configuration status is not significantly influenced by gene length. While *trans*-abundant genes are on average longer than *cis*-abundant genes (primary transcript lengths 35,969 bp versus 25,859 bp, *P* = 3.9x10^−08^; AA sequence lengths 1,012 versus 875; *P* = 0.001), corresponding with their much larger inter-mutation genomic distances, the overall correlation between gene-based *trans* fractions and primary transcript as well as AA length was not significant. Although there is an overall tendency for genes to reside in *trans* with increasing numbers of mutations, the opposite was true for the highly diverse OR genes. These were largely *cis*-abundant, their diversity being preserved by positive selection (Bustamante et al. 2005). Taken together, configuration status on the whole is neither the result of gene length nor the number of mutations. Furthermore, significant overlaps between both *cis*- and *trans*-abundant genes and MAE genes were found. First, with regard to the specific (functional) classes they were enriched for and then with regard to the shift of allele frequency distributions in these genes towards those consistent with common variation (i.e. greater allelic age on average (Savova et al. 2016)). Thus, these genes may encode common functional variation in the populations, generating wide-spread cell-to-cell, organismal, and phenotypic diversity. Their significant enrichment with functional classes involved in cell-cell and cell-environment interactions, in all populations examined, supports their general adaptive role (Savova et al. 2016). Moreover, the mutations constituting the common phase configurations might not represent disease mutations, but instead modulate the effect of individual, rare, or disease variants on gene function. To further characterize *cis*- and *trans*-abundant genes, that is, genes with either one or two altered homologues, novel approaches to the analysis of sequence-structure-function relationships will be required. These must account for the two different homologues a diploid gene possesses, including new (*in vitro*) paradigms to both separately and jointly express and characterize the two different gene products. This should also help to elucidate the differential influence of these two gene categories in functional context.

How could we explain the phenomena described? Preliminary results pointed in principle to two major mechanisms. Firstly, *cis*-abundance, with ∼60:40 *cis/trans* ratios being observed for all types of coding variants, could arise from ancestral admixture as the common underlying mechanism. Further investigations tracing human evolutionary history and population genetic processes will be required to elucidate the apparently ancient origins of *cis*-abundance and the specific admixture processes generating it. Moreover, the observation that all populations showed approximately the same *cis/trans* ratios (with some decay of LD in AFR leading to a ∼55:45 ratio) suggests that genetic admixture between two ancient populations must have occurred before dispersal of modern humans out of Africa and their population expansion worldwide (Nielsen et al. 2017; McEvoy et al. 2011). Secondly, a significant overrepresentation of particular functional categories in the global sets of phase-sensitive, and *cis*- and *trans*-abundant, genes was observed. With ≥ 2 protein-altering mutations, these genes featured an increased mutational load. As preliminary results indicated, processes of ancient balancing selection may have contributed to their higher genetic diversity to preserve their functional flexibility as an adaptive advantage (Wu and Dunlap 2002; Sellis et al. 2011; Hoehe et al. 2014; Savova et al. 2016). Moreover, the observation that all populations had the same distributional patterns of protein-altering mutations and functional enrichment suggests that these selective processes must have occurred before ancestral admixture. Thus, this conserved, potentially functionally important ‘phase-sensitive part’ of the diploid human genome may have very ancient origins. This applies particularly also to the common pairs of co-occurring mutations characterizing *cis*-abundant genes. These could serve as ‘evolutionary signals’ that could contribute to further elucidate the evolutionary history of the phenomena described. In sum, processes of ancient selection followed by admixture may have shaped the overall picture observed.

This work represents a basis for diploid genomics, or ‘diplomics’ (Tewhey et al. 2011), a ‘dual’ view of biological processes. It identifies, and focuses on, those sets of genes, the homologues of which are potentially functionally different (Wu and Dunlap 2002), and divides those further into two categories, *cis*- and *trans*-abundant genes (i.e., diploid genes with either one, or two altered homologues). Thus, it sets the stage for a diploid biology which inherently is allelically biased at all levels: chromatin organization, epigenomes, transcriptomes including transcriptional regulation, and proteomes (Dixon et al. 2015; Leung et al. 2015). Diploid biology will need to be described and understood with reference to the two homologues of the genes and regulatory sequences an individual possesses. In this context it should be pointed out what a challenge it is to currently describe the diploid nature of human genomes in words and terms that have been shaped by working with un-phased, i.e. ‘mixed diploid’ sequences leading to the perception and interpretation of a ‘one genome world’. Towards diploid genomics, major questions to be addressed in the future are, for example, whether, and how, biology could change with *cis/trans* ratios, or how phenotype could change with configuration status, and whether *cis*-abundance could represent a general phenomenon in all diploid species.

## METHODS

### Use of 1000 Genomes (1000G) Consortium data

Haplotype data from 1,092 genomes (Abecasis et al. 2012) were downloaded from ftp.1000genomes.ebi.ac.uk/vol1/ftp/phase1/analysis_results/shapeit2_phased_haplotypes/. Phase configurations were assessed in the total of 1,092 genomes and separately in the four ancestry groups contained therein (EUR n=379, EAS n= 286, admixed AMR n = 181, AFR n = 246) and their 14 populations. Phased data were available across all 1,092 genomes, with ‘no call’ rates between 2.1 and 6% and routine use of imputation in the case of missing data. Regarding the accuracy of inferred haplotypes at common SNPs, a phasing (switch) error every 300-400 kb on average has been deduced. Specifically, exome data had a substantially higher coverage (50-100X) due to the integration of additional, targeted deep exome sequence data (for all details see Abecasis et al. (2012).

### Use of 184 experimentally haplotype-resolved genomes from the Personal Genome Project (PGP) database

Individuals were recruited as part of PGP (Ball et al. 2012). Participants enrolled in the PGP gave full consent to have their genotypic and phenotypic data made freely and publically available. Documents reviewed and signed by each participant can be found at http://www.personalgenomes.org/harvard/sign-up. Each participant provided a blood sample, self-reported ethnicity, and detailed phenotype information. High molecular weight DNA was isolated, as previously described (Peters et al. 2012), for haplotype-resolved whole genome analysis using Complete Genomics’ Long Fragment Read technology (Peters et al. 2012). This enabled > 98% of all heterozygous SNPs to be placed into contigs with an average N50 of 800 kb (Mao et al. 2016). Importantly, a large number of technical replicate samples were generated to measure phasing error rates. As demonstrated previously (Peters et al. 2012), these error rates were exceedingly low (Mao et al. 2016) with ∼86% of overlapping blocks between replicate samples completely error-free. Data from these samples was projected onto a principle component analysis using four populations from the HapMap project. This confirmed that the self-reported ethnicities from each participant matched their genetic profile. Importantly, for analyses in this study, a contig filter was applied to ensure that phase configurations were determined only for those protein-altering mutations and coding variants that were contained within the same contig.

### Annotation of mutations, assessment of *cis* and *trans* configurations, evaluation of *cis/trans* ratios

RefSeq genes were downloaded from UCSC table browser (Hg19). All transcripts belonging to an autosomal gene were merged and the coordinates defining the entire gene region determined, resulting in a final set of 18,121 autosomal genes. For analysis of the 1,092 genomes, the annotation of AA exchanges, sSNPs, and mutations predicted to functionally alter the protein, was provided by the 1000G database. Annotation of AA exchanges and sSNPs in PGP data were provided by the PGP database (Mao et al. 2016). For annotation of protein-altering mutations, a combination of PolyPhen-2 (Adzhubei et al. 2010) and SIFT (Ng and Henikoff 2001) (see also Hoehe et al. 2014) as well as GERP conservation scores (Cooper et al. 2005) were applied to ensure comparability with 1000G annotation and detect all mutations that potentially alter gene function. Default threshold values for PolyPhen-2 and SIFT were used as well as GERP scores > 2.

The 1,092 genomes were filtered for genes with mutations predicted to alter the protein, and the other types of coding variants under investigation, and intermediate output files prepared: The mutations, i.e. alleles different from the reference sequence (or ‘minor alleles’), were designated ‘1’, and the alleles identical with the reference sequence ‘0’. The specific combinations of mutations representing ‘Haplotype 1’ and ‘Haplotype 2’ were contained in two adjacent columns, with the rows representing the heterozygous coordinates and their gene IDs (see also Hoehe et al. 2014). For the analysis of PGP data, we applied PolyPhen-2 and SIFT in combination with GERP (as described above) to the PGP output files generated from each of the 184 experimentally phased genomes and prepared the intermediate output files accordingly.

To assess the concrete phase configurations of all ‘phase-sensitive’ autosomal genes with ≥ 2 mutations in each of the genomes, we applied and automated the following approach: Examine column 1 representing ‘Haplotype 1’ by moving 5’ to 3’ from cell to cell, each containing allele 1 or 0 assigned to a genomic coordinate and gene ID. Remove all genes that have only one protein-altering mutation, i.e. only one cell assigned to a gene ID, to ensure that only those mutations are included that require phasing. (Upon removal of genes with one mutation, the number of different gene IDs is equivalent to the number of phase-sensitive genes in the individual genome under investigation.) Store the series of alleles across all cells (within column 1) assigned to the same gene ID as units and subject these to the assessment of phase configurations: where all stored alleles in column 1 are exclusively 1s or 0s, a *cis* configuration is scored, otherwise a *trans* configuration (see also Hoehe et al. 2014). Thus, for each genome, a result file was generated, which contained the gene IDs with an assignment of *cis* or *trans.*

This allowed immediate calculation of the *cis* fraction (%) of an individual genome as the number of genes with *cis* configurations divided by total number of genes with ≥ 2 mutations, i.e. total configuration count (equivalent to 100%), and of the *trans* fraction (%) per genome as 100% - *cis* (%). To determine the *cis/trans* ratio for totals of 1,092 or 184 genomes, or any population sample, the median values of the *cis*, and the *trans* fractions were calculated across all genomes.

To evaluate the significance of a given *cis/trans* ratio in an individual genome, we derived the composite probability of a *cis*, or *trans* configuration across all genes. We can model the probability of an observed *cis*, or *trans* configuration in a gene with *i* mutations with a Bernoulli experiment *P_i_*(*X* = 1) where *X* = 1 denotes a *cis* configuration and *X* = 0 a *trans* configuration. Thus, we have 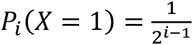 and 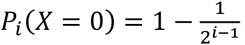. Among the number of genes with ≥ 2 mutations that have either *cis* or *trans* configurations, let *w_i_* be the relative frequency of genes with exactly *i* mutations. Thus, we have Σ_*i* ≥ 2_ *w_i_* = 1. The probability of observing a *cis* configuration among all phase-sensitive genes in a genome is then given by the weighted sum of the above defined Bernoulli probabilities: *P*(*X* = 1) = Σ_*i* ≥ 2_ *w_i_ P_i_*(*X* = 1). Thus, inserting the relative observed frequencies, *w_i_*, yielded an expected probability of ∼0.4 for a *cis* configuration to occur. The significance of the observed *cis/trans* ratio for a given genome was then computed with an exact Binomial test with *P* = 0.4. To assess the significance values for a *cis/trans* ratio which was calculated in a population sample, such as the 1,092 or 184 genomes, we derived the median values for both *cis* and *trans* configurations across all genomes, *quasi* a ‘median genome’. This was then treated as an individual genome as described above, in order to calculate a significance value for a given population sample. Thus, the significance values estimated for global *cis/trans* ratios most likely represent an underestimation.

### Simulation of phased genomes, derivation of expected *cis/trans* ratios

To verify the theoretical assumptions on the composite probability of a *cis* or *trans* configuration, simulations of phase were performed assuming that the mutations are distributed randomly between the two homologues of a gene. Accordingly, a virtual set of 1,092 phased genomes was generated as follows: for each virtual genome, random numbers of mutations were drawn in the range observed in the 1,092 genomes data set (∼2,500-3,500). The mutations were sampled from the total of ∼300,000 protein-altering mutations annotated in this data set. Phase was simulated assigning to every single mutation in a gene a 50:50 chance to exist on either homologue 1 or 2. Practically this was achieved by randomly drawing with each mutation a phase, i.e. ‘homologue 1’ or ‘homologue 2’ attached, from the 1000G database. Accordingly, a random distribution of all nsSNPs between the two homologues was simulated, drawing randomly between ∼5,500 and ∼7,500 nsSNPs with either homologue 1 or 2 attached from the entire pool of ∼1.5 Mio nsSNPs annotated in the 1,092 ‘real’ genomes, thus generating a second virtual set of 1,092 phased genomes. To test the validity of our approach to simulate phase, we assessed the *cis/trans* ratios separately for 2 up to 5 variants in both virtual data sets, and compared these ratios to the probabilities *P* for these numbers of variants to occur in *cis* under conditions of random distribution, which is 1/2^n-1^, with *n* the number of variants. Comparative evaluation showed that the *cis/trans* ratios which were simulated for defined numbers of variants were essentially identical to those expected. Thus, the simulated data were considered valid and the (composite) expected *cis/trans* ratios across all genomes in both virtual data sets were derived. The expected *cis/trans* ratios provided the basis for the estimation of the significance values of the observed *cis/trans* ratios.

The two virtual sets of 1,092 phased genomes also provided the basis to simulate the highly proportional relationships between the number of mutations per genome and the numbers of genes with 1 mutation, ≥ 2 mutations, *cis* and *trans* configurations (as mentioned in the chapter on ‘a global set of phase-sensitive genes’; see also Supplemental Table S7) as the result of a random distribution of mutations onto existing exome structure.

### Distinction of *cis*- and *trans*-abundant genes

To identify *cis* or *trans*-abundant genes, that is, genes with ≥ 2 mutations exhibiting either configuration in significant excess, we examined the ratio of *cis* to *trans* configurations for each single gene (the ‘gene-based’ *cis/trans* ratio) across all genomes in a defined population sample. Significant abundance of either configuration was evaluated with a Binomial test: Given a phase-sensitive gene, let *X* be a random variable that describes the number of *k* (out of *n*) genomes for which a *cis*-configuration consisting of two or more mutations has been found (*n* = 1,092 or any other sample size specified), we computed the corresponding *P*-value of the Binomial test with null hypothesis that *cis* and *trans* configurations were equally likely:

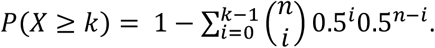

In contrast to the calculations above we used the more stringent assumption of *P* = 0.5 for a *cis* configuration as null hypothesis in order to identify *cis*-abundance for genes in the majority of genomes. Genes with *P*-value ≤ 0.05 were defined as *cis*-abundant. Likewise, genes with *P*-value ≤ 0.05 when counting the *trans* configurations were defined as *trans*-abundant.

### Over-representation analysis

In order to assess significance of over-representation of gene lists in pre-annotated gene sets (for example pathways, gene ontology (GO) terms, pre-defined gene lists, etc.) we used the hyper-geometric distribution. For each annotation set, the *P*-value is calculated as:

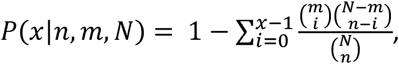

where *x* is the number of entities in the respective gene list that overlap with the entities in the annotation set, *n* is the total number of entities in the annotation set, *m* is the total number of entities in the gene list and *N* is the total number of genes (background). Since many annotation sets were tested, we routinely corrected for multiple hypothesis testing using the False Discovery Rate procedure within each type of annotation set (Benjamini and Hochberg 1995). For functional testing, we used the ConsensusPathDB tool (version 32) which holds 5,068 pre-defined pathway gene sets along with the latest GO terms (Herwig et al. 2016). Pre-defined gene lists from literature included genes that were found monoallelically (4,227 genes) and biallelically (6,006) expressed across multiple cell lines, 226 genes reported to evolve under balancing selection (BS), 104 genes with ancient derived protein-coding polymorphisms or haplotypes predating the human-Neanderthal split (HNS) and 60 genes with any evidence of human-chimpanzee *trans*-species polymorphisms or haplotypes (TSPs), as described in Savova et al. (2016). As input gene lists we have used the global set of 2,402 phase-sensitive genes as well as the global set of 7,524 genes with ≥ 1 protein-altering mutations, and, additionally, the 5,040 genes without any mutations. Furthermore, we investigated over-representation with respect to the sets of 1,227 *cis*- and 786 *trans*-abundant genes. These gene lists were computed from the 2,402 phase-sensitive genes as described above.

### Data Access

Read and mapping data for all genomes reported here are available at the database of Genotypes and Phenotypes (dbGaP) under study accession number phs000905.v1.p1. In addition, the full data package minus reads and mappings are accessible through GigaDB as part of Mao et al. (2016).

## Acknowledgments

We are most grateful to M. Vingron for critical, continued support of this work. We would like to thank H. Lehrach, A.G. Clark, T.F. Wienker, B. Caffrey for most valuable suggestions; A. Gimelbrant for providing most valuable information on mono-allelically expressed genes, and A. Toth-Petroczky for comprehensive contributions to protein structural analysis. We would also like to thank the 1000 Genomes Project Consortium.

## Authors’ contributions

M.R.H. conceived, conducted and supervised the study and wrote the manuscript. R.H. performed data analysis and provided critical input. B.A.P., Q.M., and R.D. generated the experimentally phased haplotype data. G.M.C. provided critical input. T.H. performed the bulk analysis of data. All authors provided comments and input to the manuscript.

